# The membrane microenvironment regulates the sequential attachment of bacteria to host cells

**DOI:** 10.1101/2020.11.25.397950

**Authors:** Xavier Pierrat, Jeremy P.H. Wong, Zainebe Al-Mayyah, Alexandre Persat

**Affiliations:** Institute of Bioengineering and Global Health Institute, School of Life Sciences, Ecole Polytechnique Fédérale de Lausanne, CH-1015 Lausanne, Switzerland

## Abstract

Pathogen attachment to host tissue is critical in the progress of many infections. Bacteria use adhesion *in vivo* to promote colonization and regulate the deployment of contact-dependent virulence traits. To specifically target host cells, they decorate themselves with adhesins, proteins that bind to mammalian cell surface receptors. One common assumption is that adhesin-receptor interactions entirely govern bacterial attachment. However, how adhesins engage with their receptors in an *in vivo*-like context remains unclear, in particular under the influence of a heterogeneous mechanical microenvironment. We here investigate the biophysical processes governing bacterial adhesion to host cells using a tunable adhesin-receptor system. By dynamically visualizing attachment, we found that bacterial adhesion to host cell surface, unlike adhesion to inert surfaces, involves two consecutive steps. Bacteria initially attach to their host without engaging adhesins. This step lasts about one minute during which bacteria can easily detach. We found that at this stage, the glycocalyx, a layer of glycosylated proteins and lipids, shields the host cell by keeping adhesin away from their receptor ligand. In a second step, adhesins engage with their target receptors to strengthen attachment for minutes to hours. The active properties of the membrane, endowed by the actin cytoskeleton, strengthen specific adhesion. Altogether, our results demonstrate that adhesin-ligand binding is not the sole regulator of bacterial adhesion. In fact, the host cell’s mechanical microenvironment relatively strongly mediated host-bacteria physical interactions, thereby playing an essential role in the onset of infection.

## Introduction

In the wild, bacteria predominantly live associated with surfaces. Their sessile lifestyle confers fitness advantages such as protection from predation and improved access to nutrients^1^. In the context of host colonization, the transition between planktonic and sessile lifestyles plays a functional role in mediating host-microbe interactions. Indeed, attachment to host tissue, more specifically to cells, is often a critical first step towards infection or commensalism^2,3^. As a result, the dynamics of attachment of single bacteria to host cells can dramatically influence the outcome of infection or regulate host-microbiota homeostasis.

Bacterial adhesion to abiotic materials greatly contributes to biofouling and contamination of indwelling medical device. Multiple physicochemical properties of the surface mediate adhesion to an inert substrate, including charge, hydrophobicity and conditioning^4^. In addition, mechanical properties of the material such as stiffness and surrounding fluid flow regulate attachment strength and dynamics^5–7^. This understanding of adhesion to abiotic materials provides us with rudimentary insights on adhesion to biological tissue. However, the physical and biological complexity of biotic surfaces remains overlooked when making the analogy between living and inert materials. The surface of host mammalian cells is composed of a soft lipid bilayer densely packed with surface proteins^8^. In addition, it is an active surface, permanently rearranging itself under the action of force-generating structures such as the cytoskeleton. Finally, in contrast with abiotic adhesion, bacterial attachment to host cell involves specific molecular interactions^3^. As a result, the analogy between biotic and abiotic adhesion may overlook critical physical and biological regulators.

Pathogens and commensals alike express at their surfaces proteins that specifically bind to host membrane ligands. These cell type-specific adhesins promote tissue tropism during infection or colonization^9^. These can be classified in categories that reflect their structure and molecular mechanism of display. Adhesins from the autotransporter family are exposed immediately near the bacterial cell envelope^10^. Their structure includes an outer-membrane beta-barrel scaffold and an inner alpha helix that holds a passenger domain. This domain often includes its ligand-binding domain^11^. Intimin is an autotransporter adhesin from enteropathogenic and enterohemorrhagic *Escherichia coli* that mediates attachment to gut epithelial cells. Intimin binds to Tir host membrane receptors that have been preemptively translocated by the bacterium^12^. Similarly, *Yersinia pseudotuberculosis* uses invasin, which binds to beta integrins present at the host cell membrane, to initiate host cell entry during infection^13^.

Little is known about how the host microenvironment mediates the interaction between adhesins and their receptors. Measurement of bacterial internalization suggest that the membrane fluidity of host cells slightly improves bacterial adhesion^14^. At the molecular level of single adhesins, force spectroscopy measurements have helped characterize bond mechanics both on abiotic materials and on live cells^15^. These have helped precisely identify exotic adhesin behavior such as the formation of catch bonds, which strengthen under an applied tensile force. The fimbriae tip adhesin FimH notoriously forms catch bond, allowing uropathogenic *E. coli* to strengthen adhesion in the urinary tract under flow^16–18^. Studies of adhesion, including catch bonds, have vastly focused on detachment of bacteria, where adhesion force balances externally applied mechanical load^19^. However, by focusing on the behavior of adhesins, most studies have overlooked the dynamic and physical regulation of bacterial attachment to mammalian cell surfaces.

The structure and biochemistry of adhesin-receptor interactions has been intensively characterized for a subset of adhesins^2,3,20^. One common resulting hypothesis is that the attachment behavior of single bacteria to their target host cell entirely reflects the molecular adhesin-receptor kinetics and affinity^2,9^. Does this assumption hold true when considering that adhesin and receptor must come together in a complex biophysical microenvironment? To answer this question, we combined synthetic and biophysical approaches to investigate bacterial adhesion to host cells. We engineered autotransporters for heterologous display of a synthetic adhesin on a non-pathogenic strain of *E. coli*^21^. We found that the specific attachment of bacteria to host cells occurs in two consecutive steps. A first step is unspecific, taking place within the first few seconds following contact. This is followed by the onset of specific adhesion yielding near irreversible attachment on longer timescale. We found that biomechanical properties of the host cell, including membrane rearrangement, flow and glycocalyx, regulate each of the adhesion steps. Overall, we show that the biomechanical microenvironment of host tissues strongly regulates the adhesion behavior of bacteria to their target cells, implicating that this process cannot be solely reduced to adhesin-receptor interactions.

## Results

### Synthetic adhesion to characterize bacterial attachment to host cells

To systematically probe bacterial adhesion to host cells without relying on virulence factors, we engineered an exogenous adhesin in *E. coli* and cognate receptor in HeLa cells (Fig. 1A). As adhesin, we display an anti-GFP nanobody (camelid single-domain variable heavy chain, VHH) using a truncated intimin scaffold^21,22^. The N-terminal domain consists in a beta-barrel associated with the bacterial outer membrane, through which spans an alpha helix displaying the synthetic passenger domain (Fig. S1A). Two out of three immunoglobulin-like structures of the passenger domain of wild-type intimin are replaced with an HA tag and VHH domain. We placed the construct under a tetracycline-inducible promoter. By staining with recombinant GFP and quantifying the fluorescence signal at the surface of single bacteria induced with increasing tetracycline concentrations, we generated titration curves allowing us to fine-tune the density of displayed VHH (Fig. S1D,E). To display receptor GFP ligand for the synthetic adhesin at the surface of HeLa, we displayed a doxycycline-inducible GFP fusion to a CD80 receptor anchored in the plasma membrane (Fig. S1B)^23^. Direct visualization of the fluorescence signal localized at the cell plasma membrane can confirm and help quantify receptor density (Fig. S1F).

**Figure 1:**
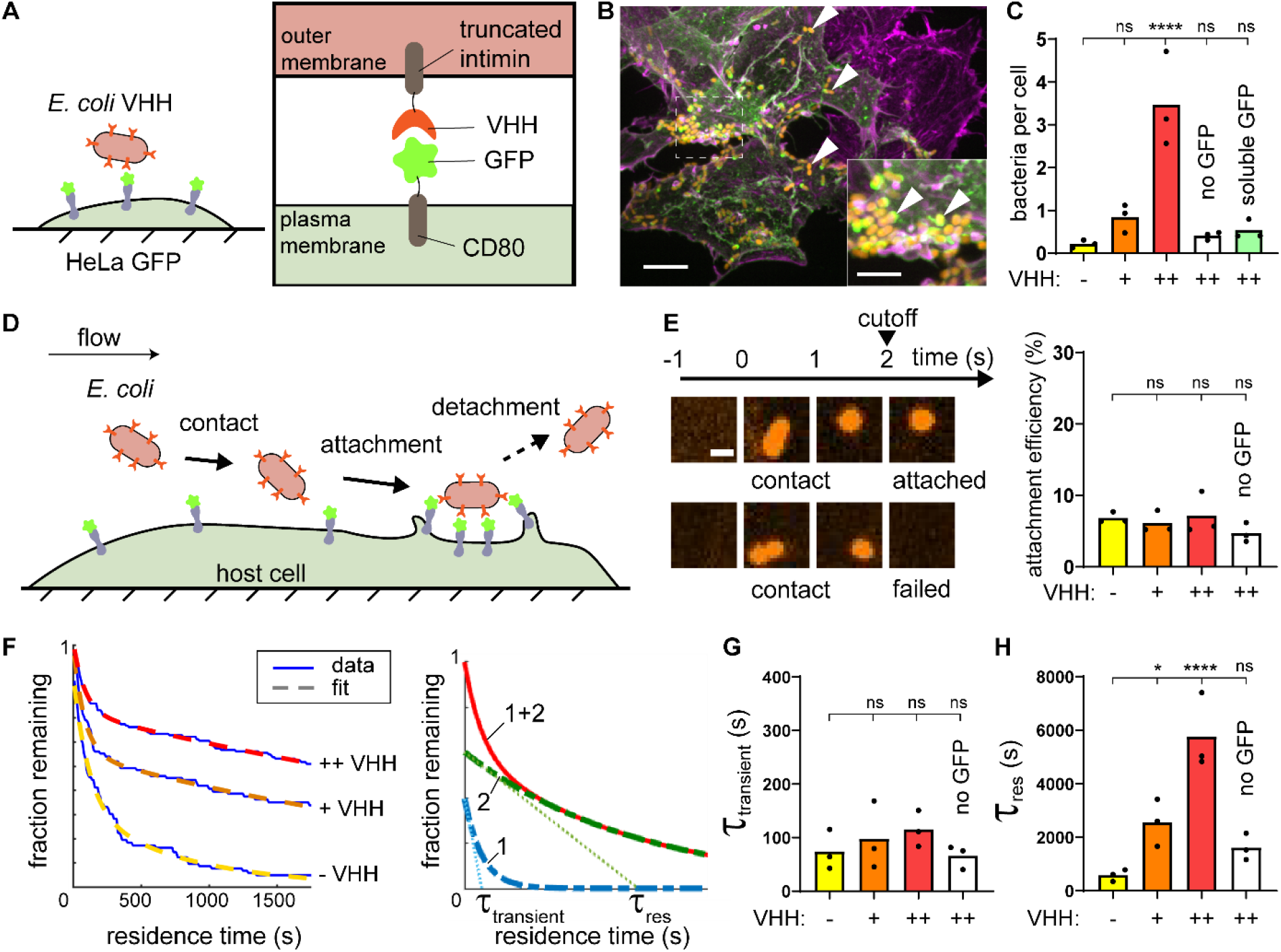
A synthetic adhesin-receptor system reveals a two-step mechanism of bacterial attachment to host cells. (A) Schematic of the synthetic adhesin-receptor system. *E. coli* cells display nanobody targeting GFP (VHH) fused to a truncated intimin autotransporter scaffold. HeLa display GFP receptors by fusion with the membrane-anchored CD80 scaffold. (B) In a mixed population of GFP+ (green) and GFP-(purple) HeLa cells, *E. coli* (orange, indicated with white arrows) specifically binds to GFP+ cells. Actin stained with phalloidin (purple). Scale bars: 10 μm (main) and 5 μm (inset). (C) Bacterial count per HeLa cell increases with *E. coli* nanobody density. *E. coli* expressing VHH at low density, or expressing VHH at high density but preincubated with soluble GFP only rarely bind to HeLa displaying GFP. (D) Dynamic visualizations of bacterial adhesion to HeLa cells under flow under flow allows to simultaneously monitor attachment and detachment events at multiple timescales. (E) Bacterial attachment efficiency is independent of VHH density and GFP display. High speed confocal imaging at 1 frame per second highlights bacterial populations that detach rapidly after contact. We considered bacteria as attached if they stayed on the HeLa surface for more than 2 s. Scale bar 2 μm. (F) We constructed residence time distributions using long timescale tracking of attached bacteria (1 h). Bare *E. coli* and *E. coli* displaying low and high VHH levels have largely different residence time distributions. We fit these distributions using the sum of two exponentials highlight two characteristic timescales *τ*_transient_ and *τ*_res_ (right illustrative graph). The single exponentials are shown in dashed green and blue and their sum is the continuous red line. (G) The model parameter *τ*_transient_ is independent of the adhesin displayed. (H) In contrast, the characteristic residence time *τ*_res_ increases with nanobody density. Statistical tests: one-way ANOVA followed by Dunnett’s post hoc test (**** P<10^−4^, * P<0.05).

We transiently transfected HeLa displaying CD80-anchored GFP, leading to a heterogeneous population of GFP positive and negative cells. We then mixed in *E. coli* with a high surface density of VHH (*E. coli* VHH) with HeLa GFP whose respective adhesin and receptor where induced separately. After washing, we visualized the co-culture by confocal microscopy. We observed that bacteria bound to GFP-positive HeLa, but not to GFP-negative cells (Fig. 1B). This indicated that the synthetic system is specific, validating it as a model of bacterial adhesion. As a result, we generated a stable and clonal doxycycline-inducible HeLa GFP-display cell line (HeLa GFP) and grew cultures of this line in microchannels to investigate adhesion in flow conditions. We diluted bacteria in mammalian cell culture medium and loaded them on a syringe pump for flow control. We injected the bacterial suspension in the microchannel covered with Hela GFP. After one hour under moderate flow, we imaged cells in the channel by confocal microscopy and quantified the number of bacteria per mammalian cell. The bacterial counts per HeLa GFP cell was larger when both constructs were induced compared to uninduced conditions (Fig. 1C). Pre-incubation of *E. coli* VHH with recombinant soluble GFP decreased the bacterial count per HeLa GFP back to the non-induced condition (Fig. 1C). Therefore, this system yields selective and dose-dependent bacterial adhesion of VHH-displaying bacteria to GFP-displaying HeLa both in static and flow conditions. Our initial characterization overall demonstrates that in tandem, *E. coli*-VHH and HeLa-GFP represent a realistic, tunable model for specific microbial adhesion to host mammalian cells.

### Bacterial attach to host cells in two successive steps

Our initial results showed that the number of bacteria attached to host cells depends on the induction levels of both VHH adhesin and GFP receptor (Fig. 1B,C). We wondered whether this was due to changes in the number of bacteria attaching or detaching from the host cell surface (Fig. 1D). This question motivated us to inspect the dynamics of attachment to HeLa cells at the single bacterium level. We tracked attachment and detachment of single bacteria over the course of 1 h (Movie S1). These visualizations helped us identify two classes of attachment behaviors. First, a large proportion of bacteria were only visible on single frames, indicating that they were in contact with the membrane for a short time. Another population of cells stayed attached for much longer times. We were intrigued by this dichotomy in adhesion behaviors and performed multiscale imaging to characterize each step.

To inspect short timescale attachment events, we performed fast confocal imaging of attachment (1 frame per second). We found that a large proportion of bacteria only stayed on the membrane for about two seconds (one or two frames, Movie S2). We then quantified the proportion of bacteria that attached to the host surface for more than two seconds relative to the total number of contacts, which we call attachment efficiency (Fig. 1E). We found that the attachment efficiency was in average only 7% when both VHH and GFP were induced. We then compared this attachment efficiency between adhesin-receptor conditions. Surprisingly, we found that neither the presence of VHH adhesins nor of GFP receptors influenced the attachment efficiency (Fig. 1E). This suggests that this early stage is not specific.

We thus speculated that the adhesin-receptor interactions regulate bacterial attachment on a longer timescale. To test this hypothesis, we timed single bacteria residing the surface of host cells during a 1 hour-long movie (Movie S1). We thus built inverse cumulative residence time distributions (Fig. 1F). We found that these distributions had exponential-like decays, which we could fit to the sum of two exponential functions (Fig. 1F and material and methods). This highlighted two characteristic timescales over which bacteria detached from the surface. The shortest timescale is on the order of 100 seconds, and was nearly identical between conditions (Fig. 1G). The longest timescale *τ*_res_, associated with the second exponential, showed large variations between VHH or GFP configurations (Fig. 1H). We measured a 10-fold increase in *τ*_res_ when bacteria displayed a high VHH density compared to bacteria displaying an empty intimin scaffold (no VHH). In addition, we measured a 3.5-fold decrease when we did not induce GFP on HeLa cells. These results implicate that adhesin-receptor interactions only materialize over minutes. As a comparison to typical association rates, we estimated the on- and off-rates of adhesin-ligand based on known kinetics constants of VHH-GFP^24^. Interestingly, the off-rate of VHH reflects a characteristic time of 6900 s, which is of the same order of magnitude as our *τ*_res_ measurements. For an arbitrary GFP concentration of 1 μM, the on-rate yields a reaction time on the order of 1 s, two order of magnitude shorter than our measurements. This suggests that other factors mediate the first adhesion step, before adhesins engage with their ligand. In summary, we highlighted that bacteria specifically attach to host cells by going through an initial non-specific attachment followed by adhesin-receptor docking, thereby promoting long lasting physical contact.

We then tested the contributions of biochemical properties of the adhesin in regulating attachment. We swapped the adhesin to two other VHH sequences coding for anti-GFP nanobodies of different affinities (*K*_D_) and kinetic rates (k_on_ and k_off_)^25^. We checked that their expression levels were unaffected using anti-HA FITC-labeled antibodies (Fig. S2i). We first verified that the fusion to intimin did not affect *K*_D_. Titrating these alternate VHH forms on *E. coli* with GFP yielded *K*_D_ matching their *in vitro* measurements performed with soluble recombinant proteins (Fig. S2ii,iii)^24,25^. We thus performed adhesion experiments on HeLa GFP under flow with *E. coli* expressing the alternate VHH forms. We observed a slight positive correlation between bacterial load per HeLa and VHH affinity across three orders of magnitude of *K*_D_ and two order of magnitude of k_off_ (Fig. S2 and S3A). Consistent with its non-specific nature, the attachment efficiency was independent of the affinity of the nanobody to GFP (Fig. S3B). On the longer timescale, we measured higher *τ*_res_ and a statistically significant increase in the pre-exponential factor C_res_ at higher affinities (Fig. S3C,D), explaining the differences in the bacterial load. Altogether, the dependence of the specific adhesion step on adhesin biochemistry was surprisingly weak compared to the changes induced by adhesin expression levels (Fig. 1H and Fig. S3C).

### Bacteria attach to abiotic surfaces in a single specific step

We suspected that the complex of physical microenvironment the host cell membrane plays a role in either of the two successive steps of attachment. To provide additional insights on these factors, we compared the specific adhesion of *E. coli* to the surface of an abiotic material with the one on mammalian cells (Fig. 2A). We engineered specific adhesion to glass by conjugating receptors to a coverslip substrate. We conjugated N-terminally His-tagged recombinant GFP to nitrilotriacetic acid (Ni-NTA) functionalized glass, on which we bonded elastomeric microfluidic channels (see material and methods). We monitored the dynamics of specific adhesion to abiotic surface by flowing a bacterial suspension in the GFP-coated microchannel. We observed bacteria almost exclusively attaching to the GFP-coated areas, thereby validating adhesion specificity (Fig. S4A-C and Movie S3). These experiments highlighted a blatant difference with mammalian cells: there were ten times more bacteria attached to the GFP-coated glass surface than on HeLa cells (Fig. 2B). This difference was strictly dependent on VHH-GFP interactions as bacteria only sparsely attached to untreated glass, or to glass coated with mKate2, a red fluorescent protein that does not bind VHH (Fig. S4D).

**Figure 2:**
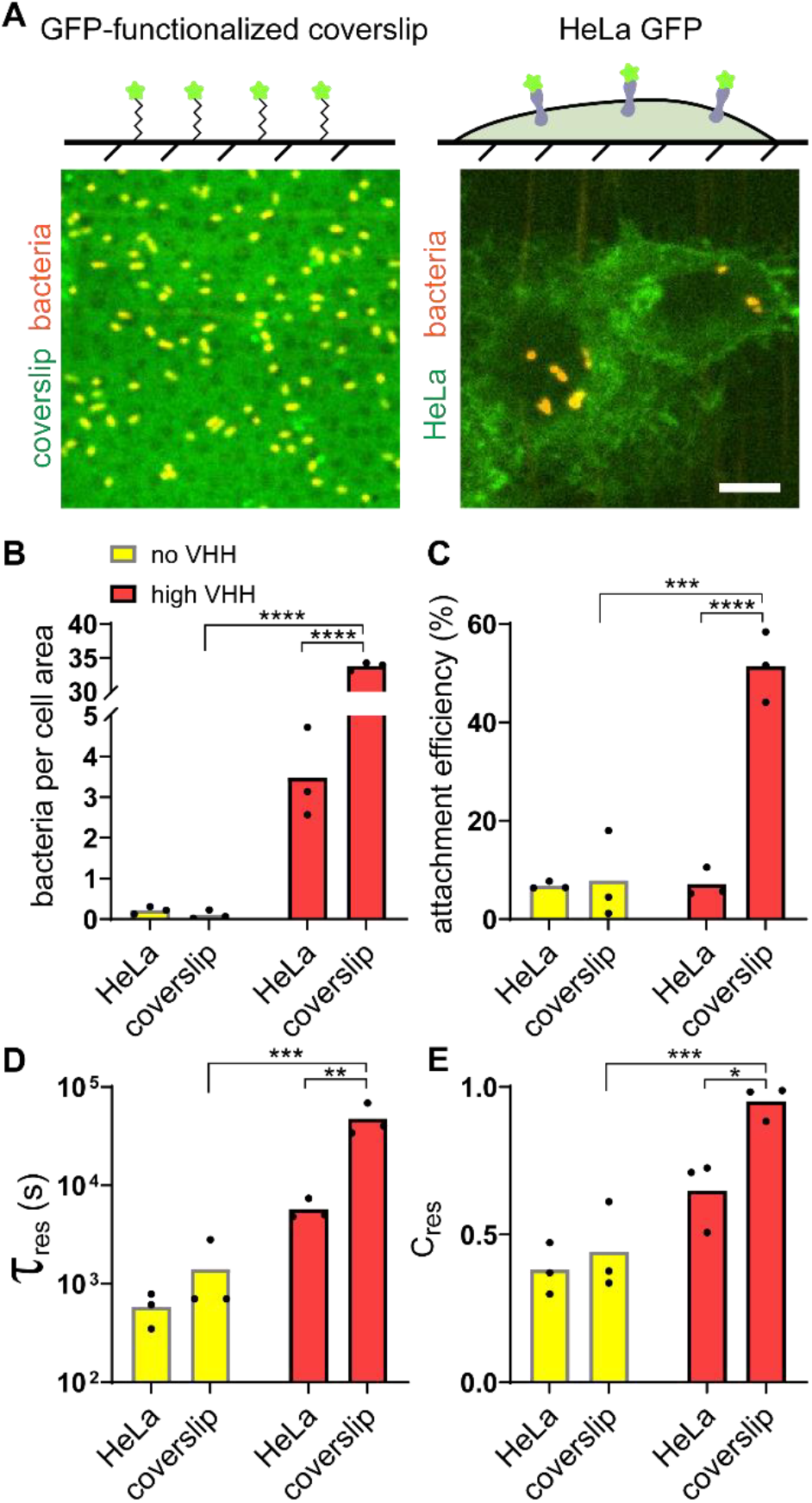
Attachment of bacteria to abiotic surface is a single step process. (A) (Top) Controlled GFP-functionalized coverslips permits visualization of specific adhesion to hard, abiotic surface and quantitative comparison with adhesion to mammalian cells. (Bottom) Representative confocal microscopy images of bacterial binding to GFP-coated coverslips (left) and HeLa-GFP (right). Scale bar: 10 μm. (B) Final bacterial count per cell area is about 10-fold larger on GFP-coated coverslips than HeLa in the presence of VHH. (C) Bacterial attachment efficiency is higher on GFP-coated coverslips than HeLa in the presence of VHH. (D) The characteristic residence time *τ*_res_ shows the VHH-dependent binding to coverslips is stronger than to HeLa. (E) Relative contribution of short and long timescale exponential fits shows that 95% of *E. coli* VHH strongly bind to GFP-coated coverslips. Statistical tests: two-way ANOVA and Sidak post-hoc test (**** P<10^−4^, *** P<0.001, ** P<0.01, * P<0.05).

To further characterize the pronounced difference in adhesion between abiotic and biotic surfaces, we focused on attachment/detachment dynamics. We compared the early attachment efficiencies and residence times of bacteria on glass with the ones on HeLa cells. First, we found that about 50% of *E. coli* VHH stayed attached to the GFP-coated glass surface upon initial contact, in contrast with the 7% of bacteria remaining on HeLa GFP cells (Fig. 2C). This largely contributed to the differences in bacterial accumulation at the end of the experiment. In addition, the characteristic residence time of *E. coli* VHH on glass was more than eight times longer than on HeLa (Fig. 2D). This characteristic time was also much longer than the duration of our visualizations so that most bacteria can be considered irreversibly attached to glass. Finally, on the longer timescale, very few bacteria transiently bound to coverslips, as highlighted by the relative contribution of *τ*_transient_ (Fig. 2E). This further supports a scenario where adhesin and receptor engage rapidly and efficiently when an abiotic surface supports the receptors.

In summary, specific adhesion to an abiotic surface is controlled by early attachment events within the first few seconds of surface encounter, consistent with *in vitro* reaction rates. Successful attachment beyond this step leads to nearly irreversible surface association. Thus, a single specific step mediates attachment on abiotic surfaces, while phenomena at both short and long timescales regulate specific attachment to host cells.

### Host cell membrane mechanics regulate bacterial adhesion

Given the differences in material properties between inert and living substrates, we hypothesized that the mechanical microenvironment of host cells may play a key role in regulating attachment. Following this intuition, we investigated the role of cells mechanics in the process of adhesion to host cells. Host cell mechanics depend on the intrinsic membrane bilayer properties but also on emergent properties provided by the actin cytoskeleton.

We observed that bacteria attached to HeLa accumulate GFP at their surface, as if they were embedded into membrane invaginations (Fig. 3Ai-ii). Given the role of the cytoskeleton in the shape and mechanics of eukaryotic cell membranes, we hypothesized that actin could play a role in bacterial attachment. To first explore this possibility, we visualized the actin cytoskeleton of bacteria-bound cells using fluorescent phalloidin staining. The actin density increased around individual attached bacteria, indicating a potential morphological remodeling of the membrane upon attachment (Fig. 3Aiii-iv). Our GFP display construct is based on a truncation of the CD80 receptor that is overexpressed in macrophages with notoriously increased actin remodeling. To exclude the possibility that remodeling is an artefact of the C-terminal CD80 anchor, we fused GFP to a glycosylphosphatidylinositol (GPI) membrane anchor devoid of cytosolic signaling components (Fig. S1C)^26^. There, we could also observe a similar actin remodeling and membrane surrounding bacteria (Fig. 3B and Fig S5A,B). The membrane remodeling occurred within minutes, on a similar timescale as the GFP uptake (Movie S4,5 Fig. S5C,D). Actin-dependent membrane remodeling could thus increase the contact area between bacteria and host cell, stimulating adhesion-receptor interactions and consequently increasing adhesion strength.

**Figure 3:**
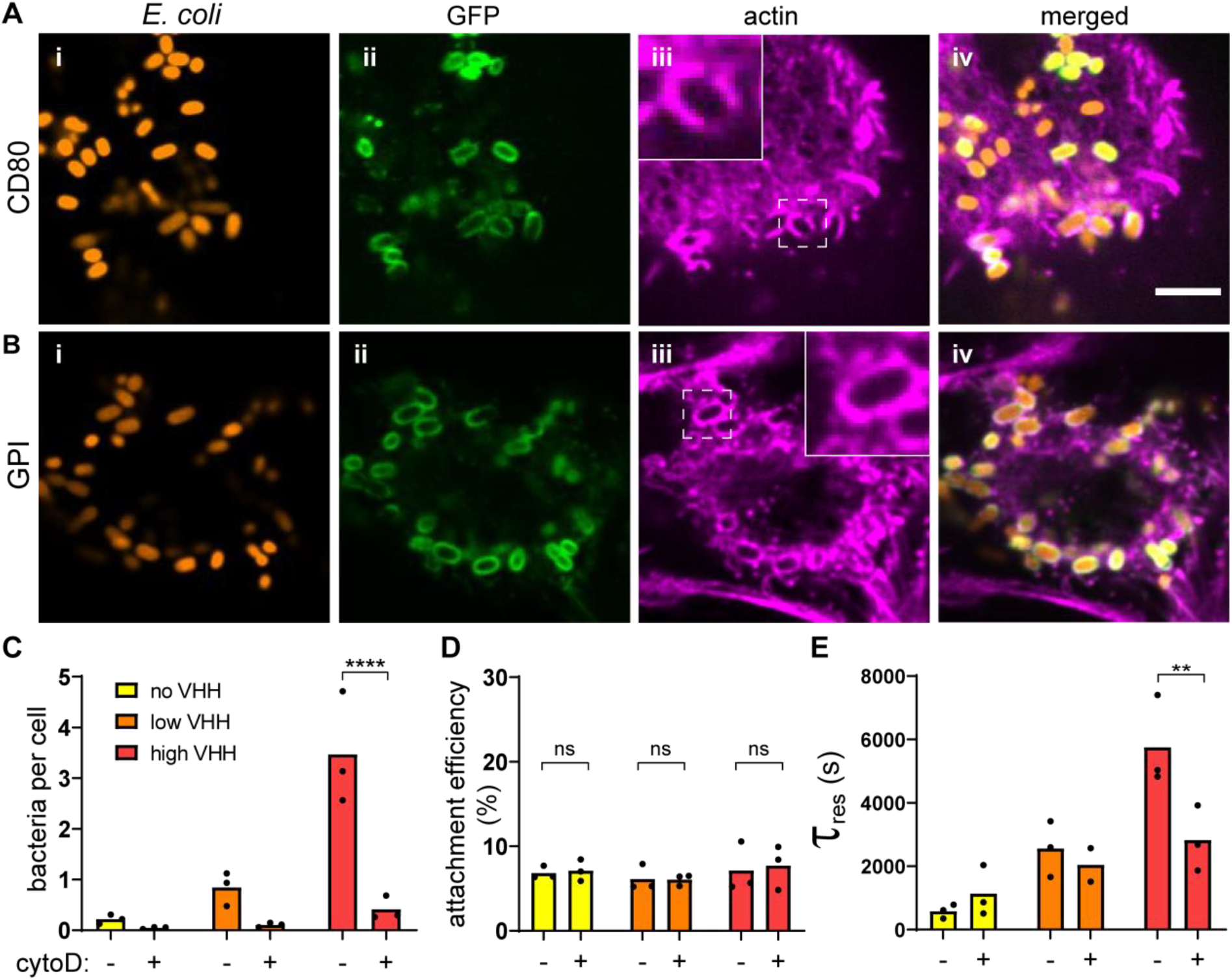
Regulation of bacterial adhesion by host cytoskeleton. (A) Actin rearranges around attached bacteria. After static incubation with *E. coli* VHH (orange), HeLa displaying GFP with a CD80 anchor (green) were stained for actin (purple). Scale bar: 5 μm. (B) Bacteria promote actin embeddings in the absence of any cytosolic component in the mammalian cell. After static co-culture with *E. coli* VHH (red), HeLa displaying GFP with a glycosylphosphatidylinositol (GPI), which does not harbor any cytosolic signaling domain, also shows strong actin remodeling around attached bacteria. (C) HeLa treatment with the actin polymerization inhibitor cytochalasin D (cytoD) reduces the bacterial count per HeLa cells. (D) Bacterial attachment efficiency is independent of actin polymerization. (E) The characteristic residence time *τ*_res_ decreases in the presence of cytochalasin D at high VHH density. Statistical tests: two-way ANOVA and Sidak post-hoc test (**** P<10^−4^, ** P<0.01).

We further tested the role of membrane remodeling in bacterial attachment by employing cytochalasin D, a drug inhibiting actin polymerization^27^. We measured an 8-fold reduction in *E. coli* VHH attachment on treated cells compared to the untreated control (Fig. 3C). Inhibiting actin polymerization did not decrease the attachment efficiency of bacteria at early timescales (Fig. 3D). However, bacterial residence time was decreased in presence of the drug (Fig. 3E). This difference was most dramatic for higher VHH densities. This suggests that membrane remodeling upon attachment takes place on the minute timescale, thereby stabilizing adhesin-receptor interactions.

Following the dependence of attachment on cytoskeletal density, which could affect membrane stiffness at the microscale, we investigated the contributions of intrinsic membrane mechanics on attachment^14^. We used a chemical approach where we modified membrane composition by growing HeLa cells in the presence of lipids known to modulate fluidity and stiffness. We first incubated cells with saturated fatty acids (margaric acid, MA) or with polyunsaturated fatty acids (eicosapentaenoic acid, EPA), which respectively increase and decrease membrane bending stiffness^28^. We could not detect changes in any of the attachment/detachment metrics compared to a negative control (Fig. S6A-D). We separately manipulated membrane fluidity by controlling HeLa’s membrane cholesterol content. We enriched the membrane with cholesterol or depleted cholesterol with methyl-β-cyclodextrin^29^. Here we measured a slight increase of bacterial load with decreasing fluidity (Fig. S6E). However, neither attachment efficiency nor residence times where affected by fluidity (Fig. S6F-H). Altogether, our results highlight that the intrinsic membrane mechanical properties such as bending stiffness and fluidity only slightly influence adhesion, while the host cytoskeleton plays an important active function in reinforcing specific bacterial attachment.

### The glycocalyx shields the host from receptor-specific bacterial adhesion

Membrane mechanics regulate how bacteria engage in specific adhesion to hosts cell on timescale of minutes. Still, membrane mechanical properties had little effect on the non-specific adhesion step, which differed so much between glass and cells, as the attachment efficiencies upon membrane and cytoskeletal perturbations remained below 10% (Fig. 3D and Fig. S6B,F). We thus still wondered why such a small proportion of bacteria could commit to specific adhesion upon encountering the host cell surface.

We reasoned that other mechanical components of the host cell surface could play a role in limiting bacterial adhesion. We thus hypothesized that the glycocalyx, a dense layer of glycoproteins and glycolipids that decorates the surface of most mammalian cells, could limit attachment. To test this, we investigated the role of the host glycocalyx in the dynamics of bacterial adhesion. We cultured HeLa cells with a deglycosylating mix of enzymes, thereby promoting its degradation (Fig. 4A)^30,31^. We confirmed specific enzymatic activity in mammalian medium by digesting fetuin, a N- and O-glycosylated control protein (Fig. S7). We then tracked bacterial adhesion dynamics at the surface of deglycosylated HeLa cells, which showed a dramatic effect. First, there was six times more bacteria attached to deglycosylated cells compared to their native, untreated state (Fig. 4B). The bacterial density on deglycosylated cells reached values close to the ones measured on glass (Fig. 2B). We further examined the specific contributions of the glycocalyx in attachment dynamics by comparing attachment efficiency and residence time distributions to the native state. Consistent with our hypothesis, we found that bacteria remained attached twice as efficiently to deglycosylated cells compared to untreated cells in a VHH-dependent manner (Fig. 4C). Deglycosylation only slightly increased the characteristic residence time, both in the presence and absence of VHH (Fig. 4D). Altogether, our data indicates that the mammalian glycocalyx shields the host cell membrane from direct engagement of bacterial adhesins to target receptors, thereby non-specifically limiting bacterial attachment.

**Figure 4:**
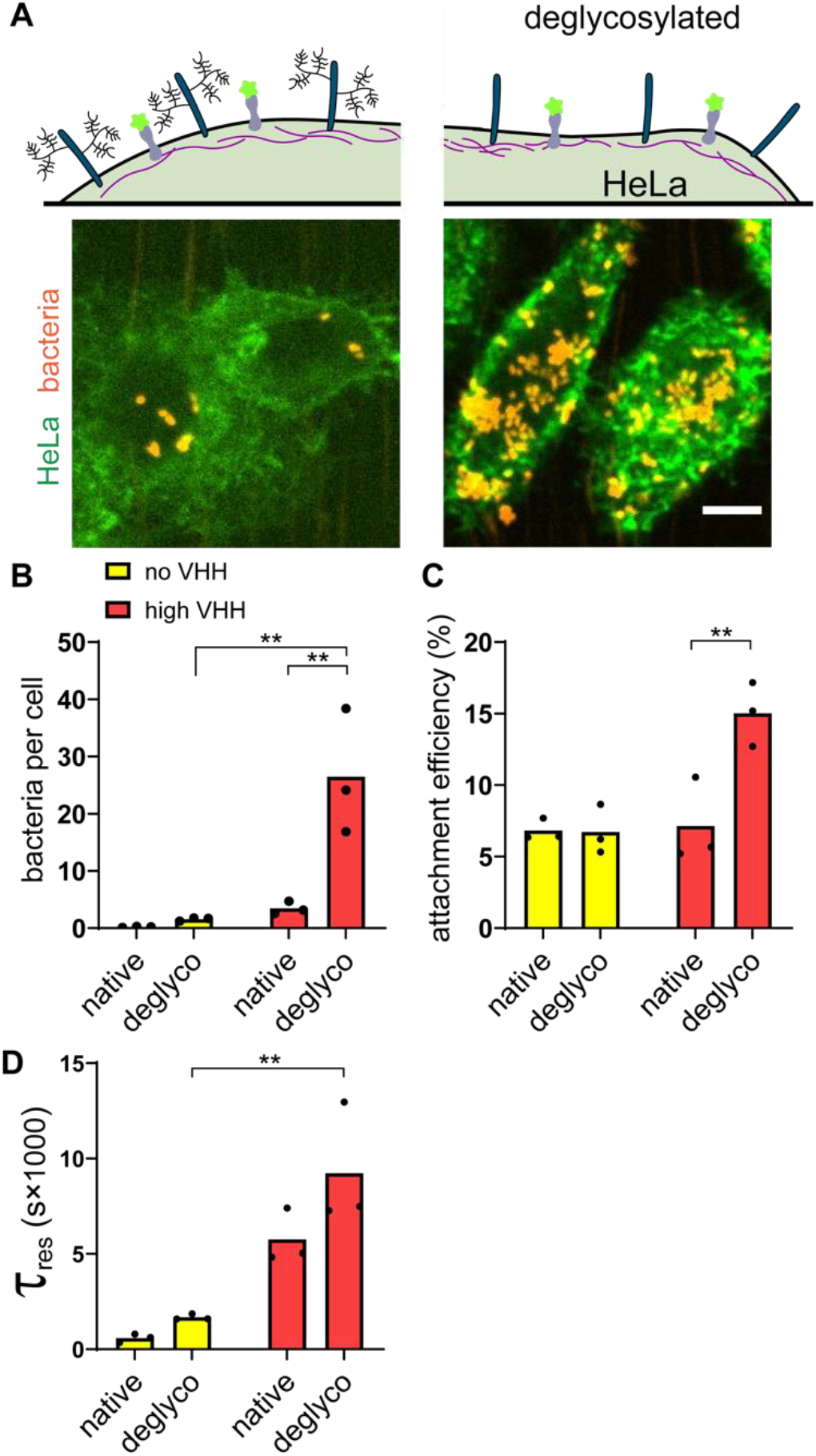
The membrane glycocalyx inhibits bacterial attachment. (A) Enzymatic deglycosylation of HeLa surface proteins increases bacterial binding. The right image shows two deglycosylated HeLa cells covered by *E. coli* VHH while the negative control in otherwise identical conditions has low bacterial count. Scale bar: 10 μm. (B-D) Comparison of bacterial adhesion dynamics between untreated cells (native) and deglycosylated cells (deglyco). (B) Final *E. coli* VHH count per HeLa cell is higher in cells lacking glycocalyx. (C) Glycocalyx removal increases the attachment efficiency of *E. coli* VHH. (D) Comparison of the characteristic residence time *τ*_res_ with or without deglycosylation mix. Statistical tests: two-way ANOVA and Sidak post-hoc test. (** P<0.01).

### Flagella and flow counteract the glycocalyx shield

Beyond simple short-range adhesins such as the ones belonging to the class of autotransporters, bacteria often display surface extensions such as flagella and fimbriae, sometimes capped with adhesins. As a result, we wondered whether such structures could help overcome the physical glycocalyx barrier by reaching through, thereby promoting the first step of adhesion. We thus explored how surface filaments could play a role in the early adhesion step. We first compared the binding of flagellated and non-flagellated bacteria to glycosylated HeLa GFP. We could not distinguish the bacterial numbers between flagellated and non flagellated strains at the end of the experiments (Fig. S8A). However, the details of attachment dynamics revealed that the flagellum mediates a tradeoff between unspecific and specific adhesion. On the one hand, we observed that flagellated *E. coli* have higher attachment efficiency (Fig. 5A). This shows that flagella promote short timescale unspecific attachment. On the other hand, the characteristic residence time of flagellated *E. coli* was more than twice shorter than its non flagellated counterpart (Fig. 5B). Consistent with this, the transient characteristic residence time was similar between conditions but the pre-exponent factor C_res_ had significantly decreased weight in our exponential fits, reflecting a high number of bacteria transiently binding and fewer bacteria strongly binding (Fig. 5C and Fig. S8B and Movie S6). Altogether, flagella mediate a tradeoff in adhesion, increasing early commitment while decreasing subsequent specific attachment.

**Figure 5:**
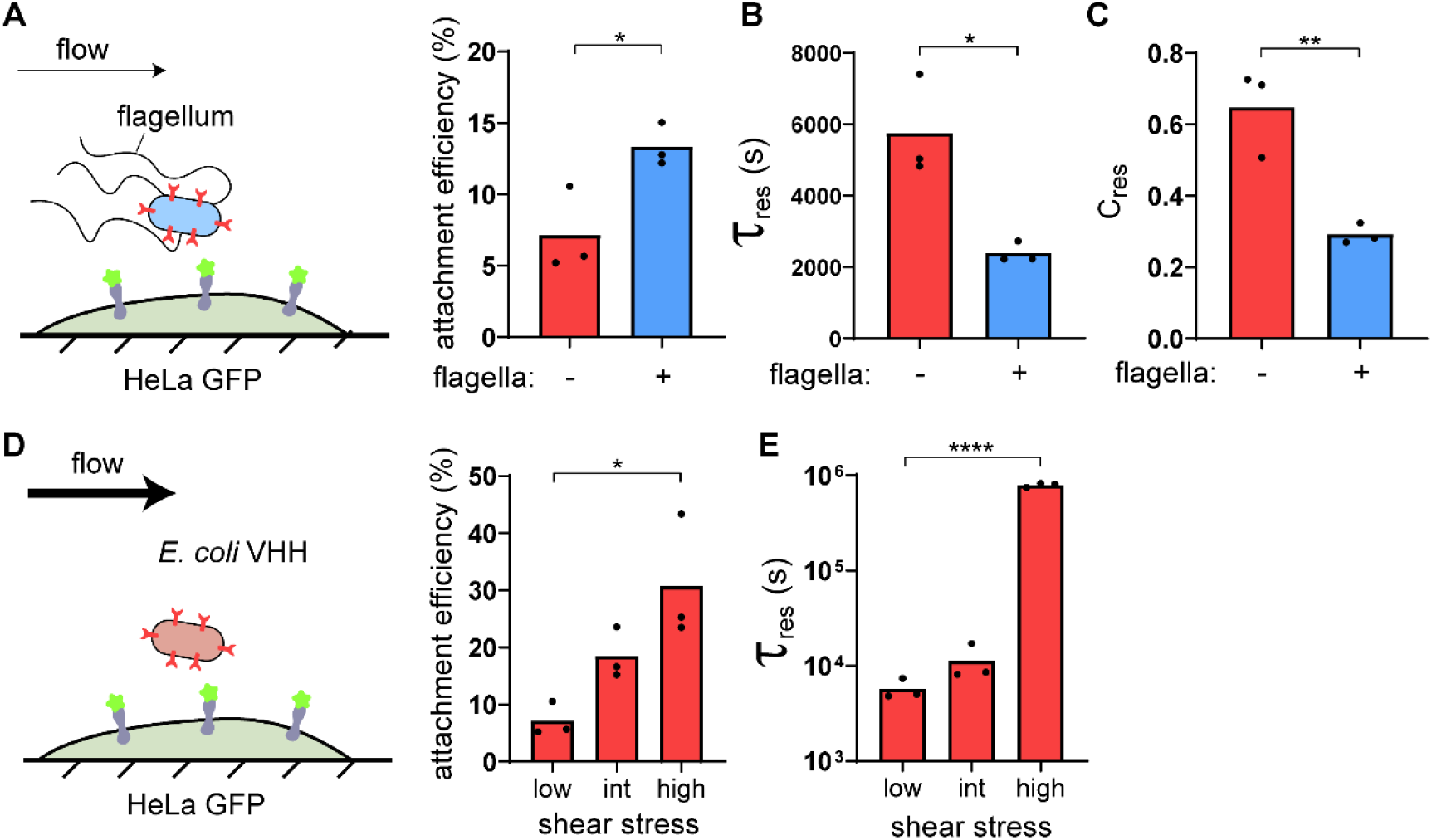
Flagella and flow attenuate the glycocalyx shield. (A) Schematic of the experimental setup. Flagellated *E. coli* VHH (blue) were compared to non-flagellated *E. coli* VHH. *E. coli* VHH attachment efficiency is increased in the presence of flagellum in flow. (B) The presence of flagella decreases the characteristic residence time in flow. (C) Comparison of the pre-exponential factor of the characteristic transient binding time *τ*_transient_ in the presence or absence of flagella shows that the proportion of bacteria strongly binding to HeLa GFP is lower with flagella. (D) We measured the attachment dynamics of *E. coli* VHH in low, intermediate (int) and high flow (respective shear stress: 0.05, 0.15 and 0.5 Pa). Bacterial attachment efficiency increases with flow intensity. (E) The characteristic residence time *τ*_res_ increases with flow intensity. In (A-C), statistical significance was calculated by two-tailed unpaired t-test (** P<0.01, * P<0.05). In (D,E) significance was calculated by one-way ANOVA followed by Dunnett’s post hoc test (* P<0.05, **** P<10^−4^).

**Figure 6:**
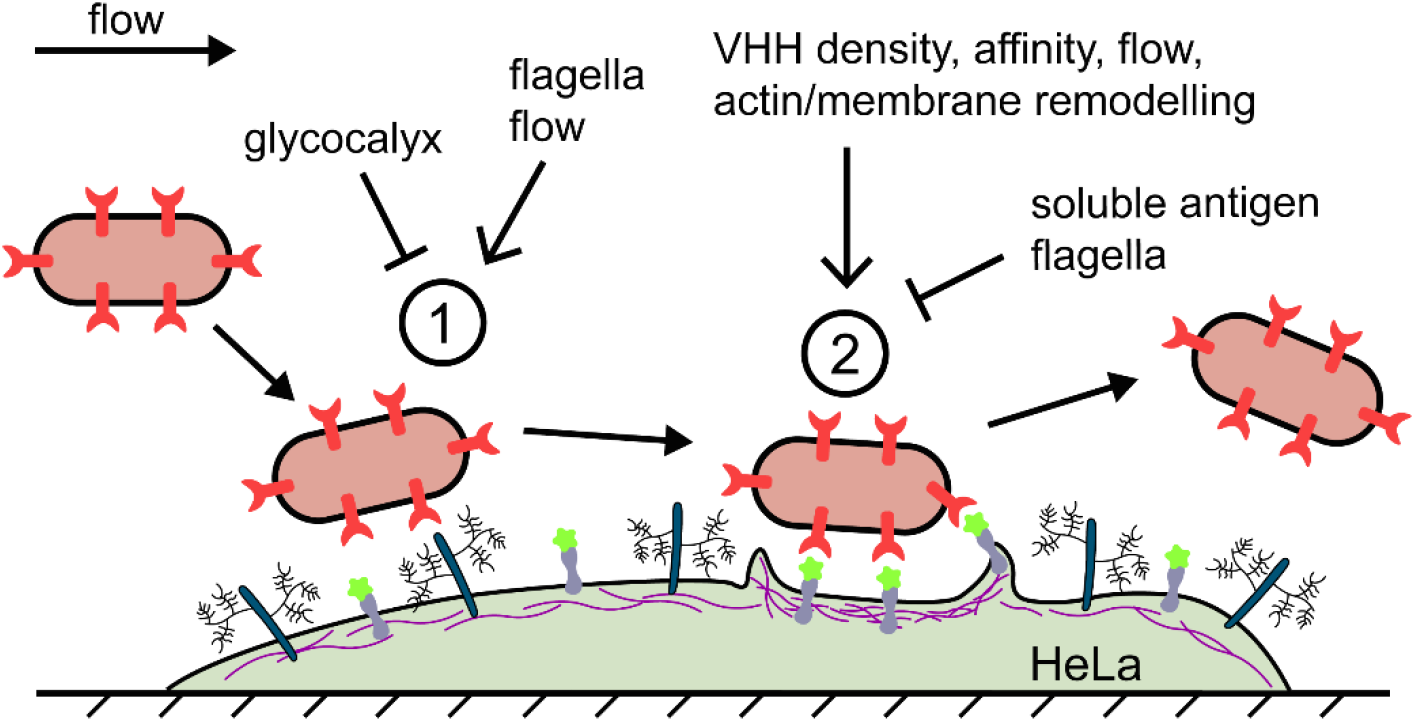
A model for mechanically-regulated, two-step bacterial attachment to host cells. Upon contact of a bacterium with a host cell, the glycocalyx blocks attachment by sterically shielding the membrane. This short timescale interaction does not involve short-range adhesins nor mammalian membrane receptors. Strong shear forces and bacterial flagellum can increase the transient binding efficiency, in part by attenuating the glycocalyx shield. The bacterium subsequently binds engages adhesins onto host receptors to promote specific adhesion. This increased adhesin density, affinity to the receptor ligand, flow, and actin polymerization promote the specific adhesion step, while the flagella and soluble antigen repress it promoting bacterial detachment.

Finally, we wondered whether fluid flow could balance the effect of the glycocalyx. Typically, hydrodynamic forces positively select for single bacteria whose adhesion force exceeds shear force. In the context of adhesion to host cells and based on molecular dynamic simulations, we suspected that flow could generate shear force that deform the ~100 nm thick glycoprotein layer, thereby reducing shielding^32^. Given these two flow-induced effects are antagonistic, we wondered how their combined contributions would ultimately affect bacterial attachment. We thus performed adhesion experiments of *E. coli* VHH to HeLa GFP at three different flow regimes. We applied flow rates that generated shear stress of 0.05, 0.15 and 0.5 Pa at the channel centerline. These stresses respectively generate 0.1, 0,3 and 1 pN hydrodynamic forces on single bacteria (assuming a bacterium is 2 μm long, 1 μm wide)^33^. We measured attachment efficiency and residence times, which are normalized metrics, that is they do not depend on the influx of bacteria in the channel.

The attachment efficiencies increased with shear stress, from 7% at low shear up to 31% at high shear. This indicates that flow promotes the unspecific adhesion within the few seconds after contact (Fig. 5D). On the timescale of minutes where adhesins engage to their GFP receptors, the characteristic residence times of bacteria increased strongly with shear stress, up to two orders of magnitude (Fig. 5E). Despite longer residence time and higher attachment efficiency in strong flow, we could not measure clear changes in absolute bacterial load per HeLa cell compared to weaker flows. We could attribute this to an unexpected decrease in the absolute number of bacterial contacts per mammalian cells with increasing flows, indicating that bacteria are less likely to encounter the host cells membrane under strong shear (Fig. S8C,D). Altogether, our results suggest that higher flows improve bacterial attachment in two ways. First, stronger flow promotes early attachment by counteracting the glycocalyx. Second, increased flow further engages adhesins to their receptors.

## Discussion

To infect or stably colonize their hosts, bacterial pathogens and commensals attach to the surface of biological tissues^34^. Adhesins are the major ingredient of bacterial adhesion to host cells. By binding to target receptor moieties at the surface of host cells, they confer adhesion specificity. We investigated how bacteria adhere to host cells by leveraging a tunable synthetic system comprising an adhesin (VHH) and a receptor (GFP). This system had been engineered for therapeutic VHH library screening and has been applied to the study of multicellular self-organization of bacterial populations^21,22^. We here repurposed it to investigate bacterial attachment to host cells while controlling adhesin expression and binding strength without affecting host viability.

We leveraged the versatility of the VHH-GFP system to perform a careful investigation of the dynamics adhesion. We first identified an overlooked temporal aspect of bacterial attachment to host cells, where a two-step sequence leads to specific attachment. After contact, bacteria attach non-specifically to host cells for not more than a minute. Bacteria subsequently engage adhesins to their receptors on a timescale consistent with adhesin-ligand rupture kinetics, in our case for minutes to hours. Sequential adhesion to host cells contrasts with the single specific step governing adhesion to abiotic surfaces.

Then, the VHH display system allowed us to test the contributions of adhesin density and binding kinetics on attachment. While VHH density promoted specific attachment, the adhesin affinity and reaction rates ended up being a surprisingly weak regulator of attachment and detachment. This could be explained by the fact after engaging several adhesins of relatively high affinity, bacterial overall avidity rapidly predominates over the affinity of individual adhesins^35^. In contrast, we found that mechanical factors of the host environment strongly regulate each of the stages of adhesion. The host glycocalyx, a layer of glycans bound to glycolipids and surface glycoproteins, inhibits the first adhesion step by physically shielding the host membrane surface. Then, we found that the host cell actin cytoskeleton shapes the membrane around attached bacteria, thereby improving specific adhesion. Membrane embedded bacteria could thus engage VHH to additional GFP receptors, thus increasing overall adhesion strength. We propose that a passive ratchet mechanism triggers the actin-dependent membrane encapsulation of bacteria^36,37^. The limited effect of membrane fluidity and bending stiffness on attachment might be explained by the fact that at the scale of a bacterium, cell stiffness is mainly driven by the underlying cytoskeleton. Overall, cell plasticity, but not intrinsic membrane stiffness, are important regulators of bacterial adhesion.

Surprisingly, we found that fluid flow improved attachment of *E. coli* VHH to HeLa GFP, both during unspecific and specific stages of adhesion. This was unexpected because fluid flow, by virtue of the shear force it generates, tends to remove bacteria from their attachment surface^33^. By shearing the glycocalyx, flow could improve the access of the bacterium to the cell membrane, thereby increasing unspecific attachment efficiency^32^. Concerning the subsequent specific step, our observations are reminiscent of flow-enhanced adhesion as a result of the formation of catch bonds^38^. However, VHH-GFP do not form catch bonds^39^. We hypothesize that flow improves specific adhesion via an indirect mechanism. For example, shearing of a bound bacterium generates tension onto the membrane, thereby stimulating actin recruitment^40^. This in turn engages more receptors, ultimately strengthening attachment. We finally note that as shear stress increases, more stringent selection for strongly attached cells could lead to the observed enhanced attachment. As a results, we cannot rule out that shear removes loosely attached bacteria at a rate that is faster than the temporal resolution of our imaging. All things considered, we demonstrated that the dependence of bacterial attachment on flow-generated forces cannot be simply extrapolated from a physically simplified behavior of a bacterium attached to a hard inert surface.

Placing our results in the context of infection showcases the breadth of strategies bacteria may have evolve to speed up and strengthen attachment to host tissue. For examples, a subset of pathogens have evolved strategies to overcome the glycoprotein shield. *Salmonella* Typhimurium actively degrades the glycocalyx during infection, a strategy suspected to promote long lasting attachment^41^. Interestingly, bacterial adhesins in some instances target surface glycans directly rather than proteins, transforming the glycocalyx shield into the target itself^42^. We have shown flagella also favor bacterial release, potentially providing an explanation for flagellum shedding it upon attachment^43^. Also, many adhesins are associated to the tips of slender filaments displayed at bacterium surface, allowing them to reach through the glycocalyx. For example, uropathogenic *E. coli* binds to mannosylated proteins at the surface of uroepithelial cells displays using the lectin FimH^18^. We expect the early attachment of bacteria expressing these long-range adhesins to be more efficient than autotransporter-based adhesins. Consistent with this, we found that flagella had a positive effect on the early attachment efficiency, even though it does not express specific adhesins.

During the process of infection, bacteria use an arsenal of virulence factors. These are deployed in a timely fashion in response to relevant signals. Synchronizing expression of virulence factors with host cell contact could promote timely deployment^33^. For example, enteropathogenic *E. coli* transfer the intimin adhesin receptors Tir to gut epithelial cells upon contact^2^. The unspecific first step of adhesion thus offers a window of opportunity to deploy these systems within minutes.

Altogether, we have demonstrated that bacterial attachment to host cells vastly differs from the expected behavior of simple adhesin-receptor interactions. Adhesin biochemistry and the physics of adhesion to inert materials only poorly predict adhesion to mammalian cells. This has therefore important implication in our view of infection. In the current context of a concerning rise in multidrug resistant pathogens, our work provides new insights that could inspire us in developing anti-adhesive therapeutics^3^.

## Material and methods

### Cloning

Plasmid cloning strategy and primers sequences are described in supplementary tables S1 and S2, respectively. Cloning was performed by restriction enzymes (NEB) and ligation with T4 ligase (Bioconcept) or by Hi-Fi Gibson assembly (NEB). PCR were performed using Phusion polymerase (Life Technologies) and DNA purification with commercially available kits. Chemically competent XL10Gold (Agilent) were used for transformation.

### Cell culture, engineering and induction

HeLa cells were cultured in DMEM (Thermofisher) supplemented with 10% FBS (Life Technologies) at 37°C and 5% CO_2_. Prior to experiments, cells were trypsinized and resuspended in FluoroBrite (Life Technologies) supplemented with 10% FBS and 1% Glutamax (Life Technologies). Cells were seeded at 100,000 cells/mL in 96 well plates or 400,000 cells/mL microchannels (Ibidi μ-Slide VI 0.4) one day prior to experiments. In microchannels, first 30 μL of cell suspension were added. Cells were left to adhere for 5-6 hours, and then reservoirs were filled with 120 μL medium.

Unless stated otherwise, we used HeLa cells displaying a doxycycline-inducible CD80-anchored GFP. To generate a stable cell line, we produced lentiviruses in HEK293T cells. Cells at 50% confluence were co-transfected with pMD2G (Addgene 12259), pCMVR8.74 (Addgene 22036) and pXP340 (Table S1) using Lipofectamine3000 (Life Technologies). Medium was changed at day 1 and lentiviruses were collected at day 2 and 3, separated from cell debris by centrifugation, sterile filtered and added to HeLa cells. Cells were selected with G418 (Chemie Brunschwig) at 300 μg/mL and resistant clones were obtained by limiting dilution in 96 well-plates. The resulting monoclonal cell line (HeLa GFP) were induced overnight with doxycycline (HiMedia) at 300 ng/mL.

HeLa cells transiently expressing GPI-anchored were obtained by lipofection of the plasmid PeGFP_GPI.

### Bacterial culture, engineering and induction

*E.coli* K12 (BW25113) were cultured in LB at 37°C. Bacteria were stably engineered to express cytoplasmic mScarlet using pZA002 for Tn7 insertion^44^. pZA002 consists in a synthetic constitutive promoter upstream of mScarlet ligated into pGRG36 for chromosomal integration. Deletion of the flagellum was performed using the lambda red system and the PCR product using oXP851 oXP852 on *E.coli* genomic DNA to delete the *FliCDST* operon^45^ (table S2). Flagellated and aflagellated fluorescent *E.coli* were then electroporated with tetracycline-inducible intimin-based display constructs. pXP383 coding for the display of VHH of medium affinity was used in this study in aflagellated *E.coli* unless stated otherwise. pXP384 and pXP388 display the VHH of lower and higher affinities, pDSG323 the empty scaffold and selected with kanamycin (Sigma) at 50 μg/mL^22^. To prepare adhesion experiments, early stationary pre-cultures were diluted 1:3000 and induced with sublethal doses of tetracycline (Sigma, 50 or 250 ng/mL) overnight under shaking conditions.

### Cell membrane and cytoskeletal perturbation

HeLa were cultured overnight with eicosapentaenoic acid (Cayman) or margaric acid (Sigma) at 150 μM. Water-soluble cholesterol (Sigma) at 1 mg/mL or cholesterol-depleting methyl-β-cyclodextrin (Sigma) at 20 mg/mL was added for 1 hour prior to the experiment. Cytochalasin D (Sigma) at 1 μM was added 5 minutes prior to- and during the experiment.

### Attachment with soluble GFP

Soluble recombinant GFP was added to the bacterial suspension at 10 μg/mL 5 min prior to the experiments.

### Generation of a Ni-NTA functionalized glass surface for selective protein immobilization

Addition of the Ni-NTA functionality to a glass surface was inspired by existing protocols^46,47^. Glass coverslips (#1.5) were placed in a holder and sonicated in acetone for 30 min. The coverslips were then rinsed with MilliQ water, dried with a stream of nitrogen gas and plasma treated for 10 minutes at maximal power (Zepto, Diener electronic). The plasma-treated coverslips were then transferred into 150 mL of 1% (v/v) (3-Aminopropyl)triethoxysilane (APTES) (Sigma-Aldrich) in toluene (Sigma-Aldrich) and stirred for 30 min. The cover slips were then rinsed in 150 mL of toluene for 10 minutes, dried by a stream of nitrogen gas, then baked at 80°C for 45 min. The coverslips were then cooled down with a stream of nitrogen gas and transferred into a 150 mL stirred solution of 2 mg/ mL p-Phenylene diisothiocyanate (PDITC) (Sigma-Aldrich) in 10% (v/v) anhydrous pyridine (Sigma-Aldrich) and 90% (v/v) N,N-dimethylformamide (DMF) (Sigma-Aldrich) for 2 h in darkness. The cover slips were then flushed with 1 volume of absolute ethanol, followed by a wash in acetone for 10 min and drying with a stream of nitrogen gas. Then half the cover slips were laid on a flat surface. We then prepared a solution of 457 mM N,N-Bis(carboxymethyl)-L-Lysine-hydrate (Sigma-Aldrich) in 1 M NaHCO3 (Sigma-Aldrich). 90 μL of the N,N-Bis(carboxymethyl)-L-Lysine-hydrate solution were deposited onto the cover slips, then sandwiched with another coverslip on top, These were incubated overnight at room temperature. The unreacted PDITC was then blocked by immersing the coverslips into a solution of 5mg/ml BSA + 5% ethanolamine in PBS for 30 min. The slides were then washed in 1x PBS for 10 minutes under constant stirring, and transferred into a solution of 1% (w/v) solution of nickel sulfate (NiSO_4_) for 1 hour under stirring, then washed in 1x PBS for 10 min followed by a second wash in 0.1x PBS for 10 minutes and dried under a stream of nitrogen gas. 50 μL of recombinant GFP protein at 1 mg/mL were deposited onto each coverslip and incubated over 2 days in the dark at 4°C. The slides were again flushed in 1x PBS for 10 minutes followed by a second wash in 0.1x PBS for 10 minutes then dried with a stream of nitrogen.

### Visualization

For widefield visualizations, we used a Nikon TiE epifluorescence microscope equipped with a Hamamatsu ORCA Flash 4 camera and an oil immersion 100x Plan APO N.A. 1.45 objective.

For all time-lapses and mammalian cell visualizations, we used a Nikon Eclipse Ti2-E inverted microscope coupled with a Yokogawa CSU W2 confocal spinning disk unit and equipped with a Prime 95B sCMOS camera (Photometrics). For time-lapses, we used a 40x objective with N.A. of 1.15 to acquire z-stacks with 2 μm intervals over 6 μm. Each plane was acquired at low laser power for 200 ms allowing to threshold out free bacteria in flow from bound bacteria. For stained mammalian cell visualizations, we used a 100x oil immersion objective with N.A. of 1.45 to acquire z-stacks with 0.5 μm intervals.

We used NIS Elements (Nikon) for three-dimensional rendering of z-stack pictures.

### Flow experiments and data acquisition

Bacteria induced overnight were diluted 1:10 in Fluorobrite 10% FBS 1% Glutamax and loaded in syringes. 3 mm/s mean flow (unless stated otherwise) was applied using syringe pumps connected to microchannels seeded with induced HeLa at 50-80% confluency. Z-stacks for bacterial attachment efficiency was generated by confocal microscopy every second. Three different fields of view were sequentially imaged for 5 min per biological replicate. Data to model residence time was generated by confocal microscopy of z-stacks every 10 s. Three different fields of view were simultaneously imaged for 60 min per biological replicate. Cell surface area was acquired once in the green channel at the start of the experiment. Number of HeLa cells was then approximated based on their average size as manually determined with 5 biological replicates of 3 frames each.

Illustrative confocal time-lapse with both channels for GFP and mScarlet were acquired at either 2 or 6 stacks per minute at 100x magnification.

### Bacteria tracking

We use the maximum intensity projection of full stacks to detect attaching bacteria. We used the Fiji plugin Trackmate with LoG detector^48^. Threshold was set so that >95% of bacteria are detected on the final frame and <5% of the tracks were false positive (two different bacteria slowing down in the same area on consecutive frames). The LAP tracker was used with 5 μm maximal inter-frame distance and gap closing, track splitting and closing with a maximal distance of 3 μm. Final number of spots, tracks and spots statistics were exported for data analysis.

### Data analysis and modeling

Data generated by trackmate was analyzed using Matlab. In brief, attachment efficiency was defined as the number of tracks strictly longer than 2 frames, divided by the total number of contacts (bacterium appearing on one frame or more). Bacteria present from the first frame were removed from the analysis to exclude bacteria that attached during handling time.

Residence times of tracks strictly longer than two frames were considered and sorted in a histogram of 10 s bins. We further transformed this data into an “inverse” cumulative histogram to present results in a manner classical for adhesion events by defining:

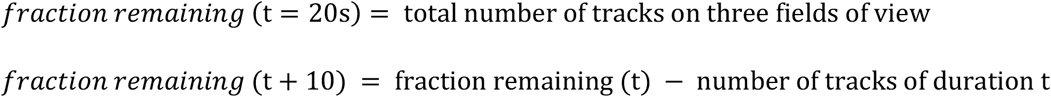

Because many bacteria were bound at the end of the acquisition, we had to circumvent the artificial stop of tracks. To do so, we considered the binding events occurring within the first 30 minutes and followed them over 30 additional minutes for the fitting. We fitted the fraction remaining as a function of residence time with a dual exponential decay as follows:

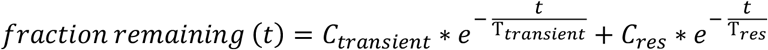

### Static co-culture and mammalian cell staining

Mammalian cells were co-incubated with bacteria for 5h 30min at a multiplicity of infection (MOI) of 50 (Fig. 1C) or for 1h at a MOI of 200 (Fig. 3A,B and S5A,B). Wells were washed once with PBS, fixed in 4% paraformaldehyde for 20 minutes, permeabilized with 0.1% Triton X-100 for 5 min and washed twice with PBS. Phalloidin-Atto 655 (Sigma) was used to stain actin at 500 nM for 15 min. DAPI was used for nuclei counterstain at 1 μM for 5 min. Cells were washed twice with PBS and imaged by confocal microscope at 100x magnification.

### Bacterial staining, titration and quantification

Bacteria displaying VHH were washed with PBS and stained with recombinant GFP at 100 μg/mL for 10 minutes prior to two PBS washes and imaging under a 1% agarose PBS pad. Wide field fluorescent pictures were taken at 100x and 1.5x lens magnification.

### Production of recombinant proteins

eGFP sequence (Genbank accession 8382257) was cloned into pET28a (Novagen) in frame with a N-terminal 6xHis tag and the resulting pXP226 was retransformed into BL21 strain. Production was induced with 1 mM IPTG (Fisher bioreagents) at 20°C overnight. Bacteria were pelleted and lysed by sonication in lysis buffer (Tris 100mM, NaCl 0.5M, glycerol 5%) and eGFP was purified using fast flow His-affinity columns (GE Healthcare) and eluted with 500 mM imidazole. Buffer was exchanged to PBS using 30kDa ultracentrigation spin columns (Merck) and aliquots at 1 mg/mL were snap frozen for further use. mKate2 was produced using the same protocol using the plasmid SpyTag003-mKate2.

## Acknowledgements

We thank Dr. Bruno Correia and Stephane Rosset at Ecole Polytechnique Fédérale de Lausanne (EPFL) for the production and purification of recombinant proteins, Dr. Ingmar Riedel-Kruse (Stanford University) for the tetracycline-inducible nanobody display constructs, Dr. Gisou Van der Goot (EPFL) for the GPI-anchored GFP construct, Dr. Didier Trono (EPFL) for the HEK293T cells and lentivectors and Dr. Joerg Huelsken (EPFL) for the CD80-GFP construct and HeLa cells. We are grateful for the fundings from the Gebert Rüf Foundation, project number GRS-057/16, the EPFL School of Life Science interdisciplinary PhD program and the Swiss National Science Foundation, project number 514495.

