## Supplementary information for "The membrane microenvironment regulates the sequential attachment of bacteria to host cells"

**This PDF file includes:**

Figures S1 to S9

Legends for movies S1 to S6

Supplementary methods

Table S1 and S2

SI references

**Other supplementary materials for this manuscript include the following:**

Movies S1 to S6

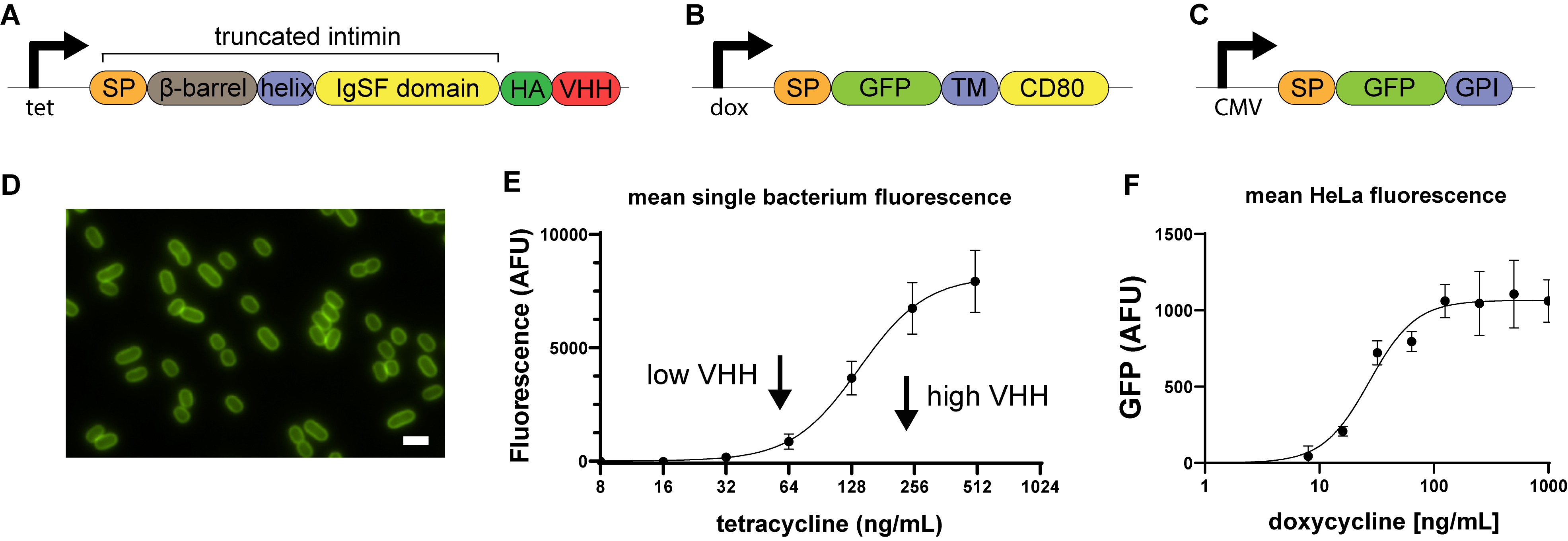

**Figure S1: Synthetic VHH adhesins and GFP receptor constructs.**

(A) Schematic of the VHH display genetic construct. *E. coli* display anti-GFP nanobodies (VHH) based on a tetracycline-inducible promoter (tet) and a truncated intimin that consists in a signal peptide (SP), a beta-barrel anchored in the outer membrane, an alpha-helix crossing the beta-barrel. The scaffold allows the display of an HA epitope tag (HA) and a nanobody anti-GFP (VHH). (B) Schematic of the GFP display construct. GFP was displayed at the surface of mammalian cells by the mean of a doxycycline-inducible promoter, a mammalian signal peptide (SP), and anchored in the plasma membrane with a CD80 transmembrane domain (TM) and its C-terminal cytosolic domain. (C) Schematic of the GFP display genetic construct devoid of cytosolic component. GFP was displayed at the surface of mammalian cells by the mean of a constitutive CMV promoter, an insulin signal peptide (SP), and anchored in the plasma membrane with a CD55 glycosylphosphatidylinositol anchor (GPI). (D) Widefield epifluorescence image of *E. coli* VHH stained with recombinant GFP. Scale bar: 2 μm. (E) Tetracycline titration followed by staining with recombinant GFP. Error bars represent standard deviation of triplicate fields of view. The solid line represents a Hill function fit. (F) Doxycycline titration on HeLa stably engineered with a doxycycline-inducible GFP display. Error bars represent standard deviation of triplicate fields. The solid line represents a Hill function fit.

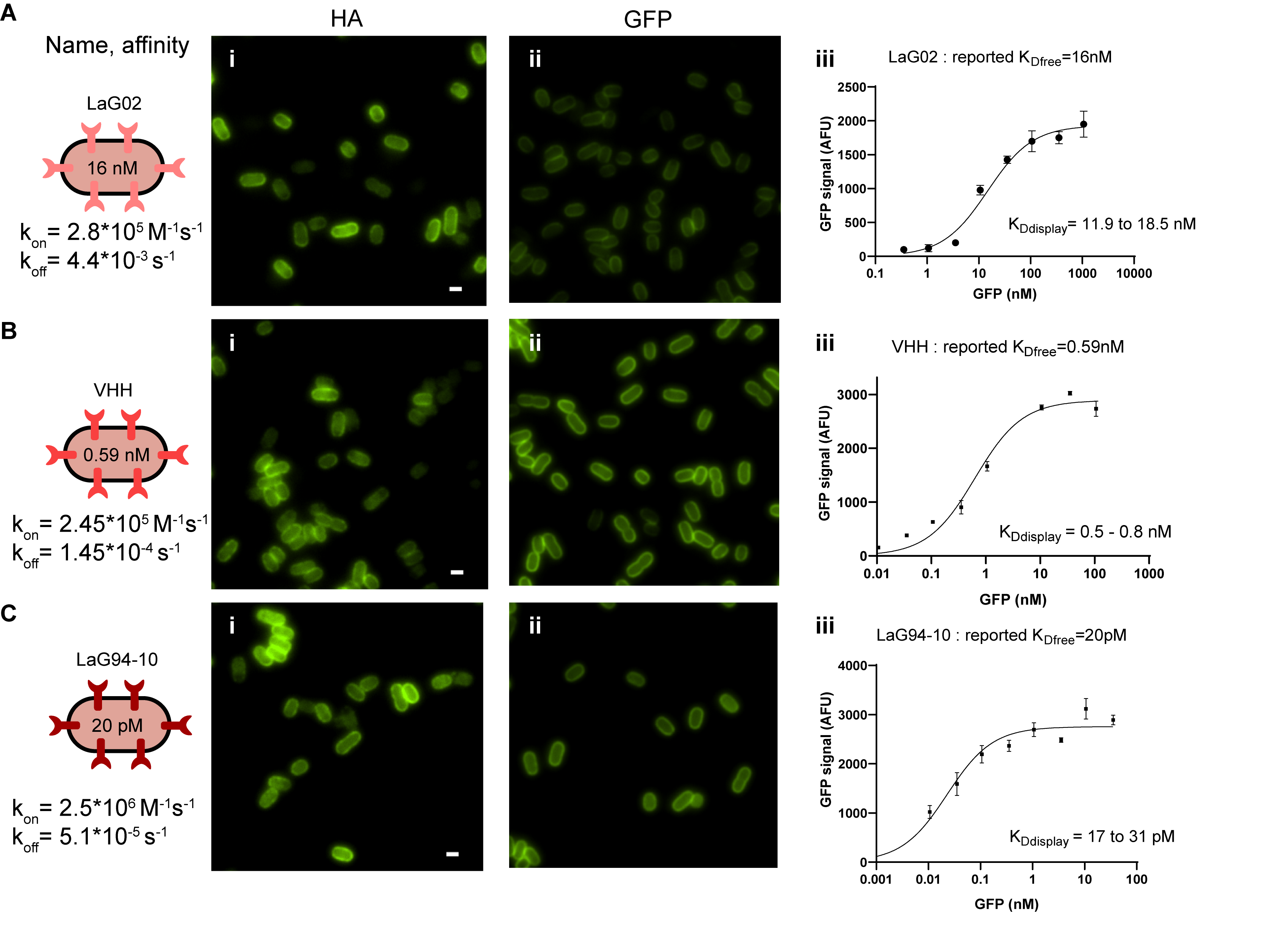

**Figure S2: Functional validation of displayed VHH based on their affinity .**

(A-C) We displayed three different anti-GFP nanobodies at the surface of *E. coli*. These have distinct biochemical properties including affinity (K_D_), on and off rates. (i) Their level of expression was measured by HA staining. Image intensity scale is identical between samples. Scale bar: 1 μm. (ii) Their functionality was then assessed by staining with excess amount of recombinant GFP: Image intensity scale is identical between samples. (iii) By titrating with recombinant GFP, we could determine the K_D_ of displayed VHH. These remain in the range of the published values obtained with soluble recombinant nanobodies. Bacteria were induced with 250 ng/mL tetracycline overnight prior to staining.

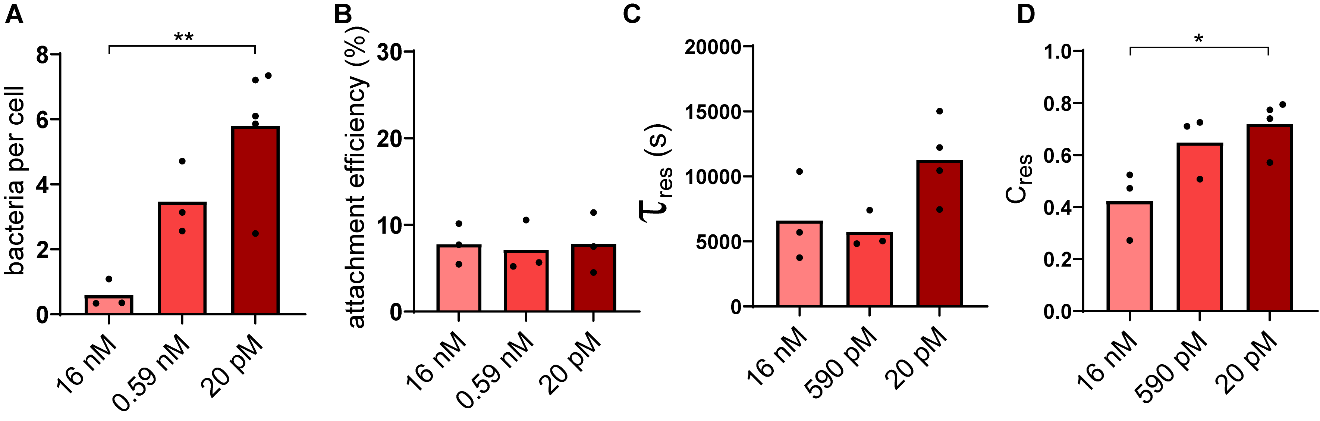

**Figure S3: Bacterial attachment as a function of adhesin affinity to GFP.**

(A) Final *E. coli*-VHH count per HeLa cell positively correlates with nanobody affinity in flow. VHH induction level is high (B) The attachment probability does not depend on the VHH affinity to GFP. (C) Comparison of the characteristic residence time 𝜏_res_ as a function of nanobody affinity. An outlier of value 775,000 s was excluded from further analysis. (D) Higher affinity increases the proportion of bacteria irreversibly binding to cells. Statistical tests: one-way ANOVA followed by Tukey post hoc test (** P<0.01, * P<0.05).

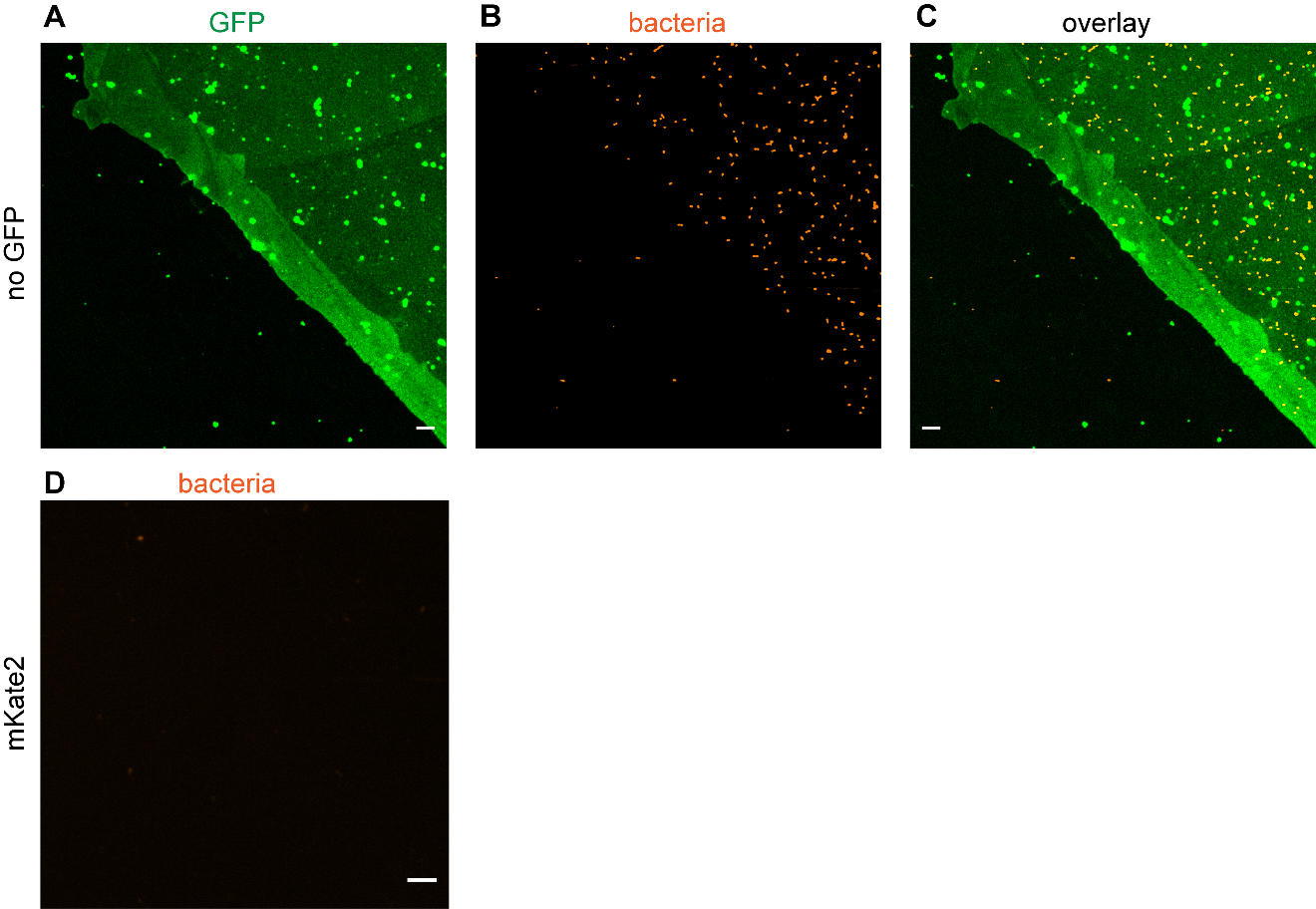

**Figure S4: Specificity of adhesion of *E. coli*-VHH to GFP-coated glass coverslips.**

(A) Microscopy image of the edge of a GFP-coated region on the coverslip. (B) Maximum intensity projection of the corresponding field showing bacterial attachment (red) after 10 min in flow. (C) Overlay of (A) and (B). (D) Maximum intensity projection of *E. coli* VHH (red) after 60 min in flow on mKate2-coated coverslips. Scale bars: 10 μm.

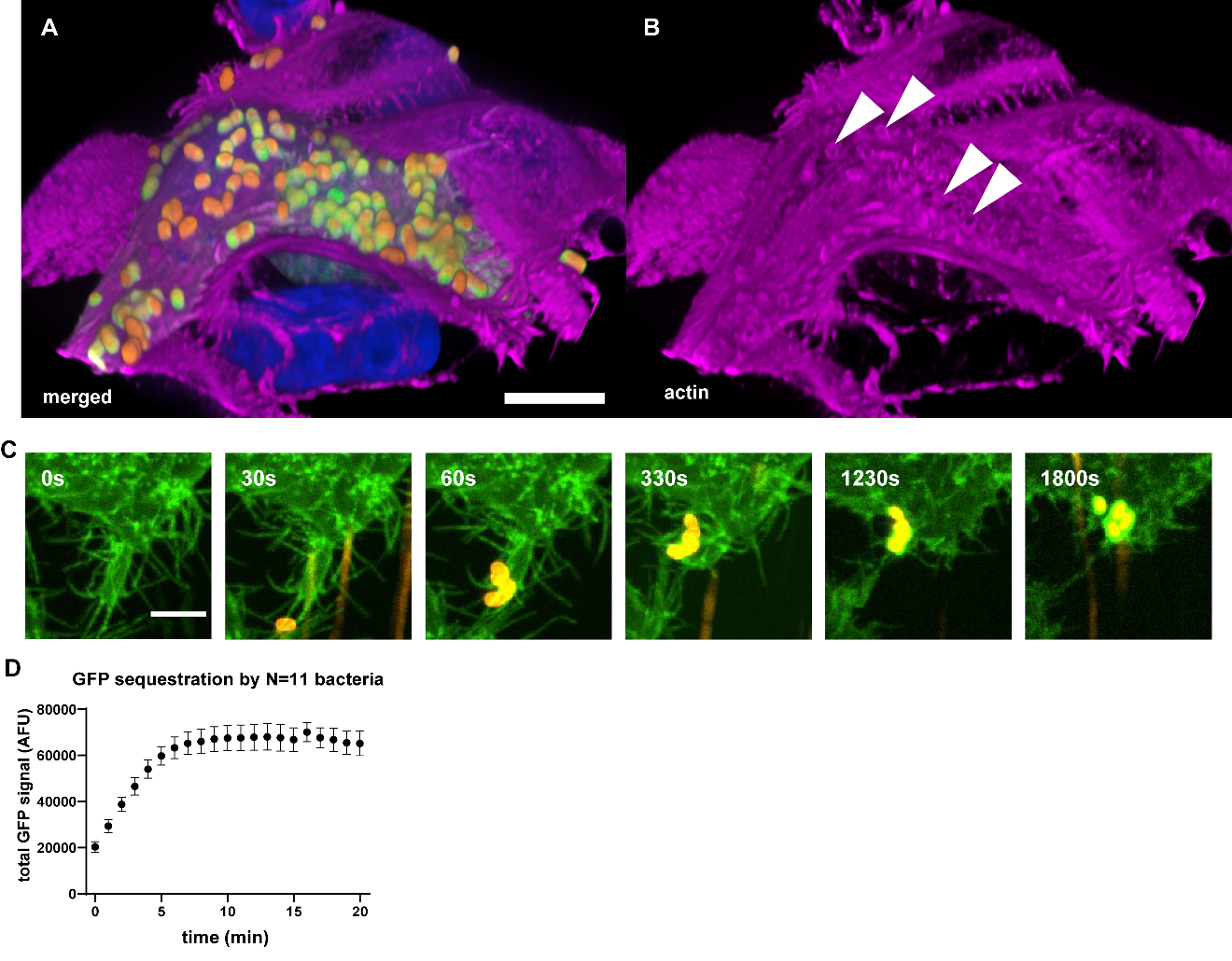

**Figure S5: HeLa cells actively remodel plasma membrane around bacteria.**

(A,B) Merged 3D visualization of a confocal Z-stack After 1h of *E. coli* VHH (red), HeLa displaying GFP with a GPI anchor (green) in static co-culture. Actin was stained with phalloidin (pink) and nuclei with DAPI (blue). Events of actin remodeling are indicated by white arrows in (B). Scale bar: 10 μm. (C) HeLa GFP (CD80) cells actively pull bacteria towards their cell body. Maximum intensity projection of a confocal time-lapse experiment of *E. coli* VHH (red), HeLa GFP (green) and flow. Corresponds to supplementary movie S4. Scale bar: 5 μm. (D) Bacteria sequester GFP within minutes in static conditions. Quantification of the total fluorescence of N = 11 bacteria in supplementary movie S5.

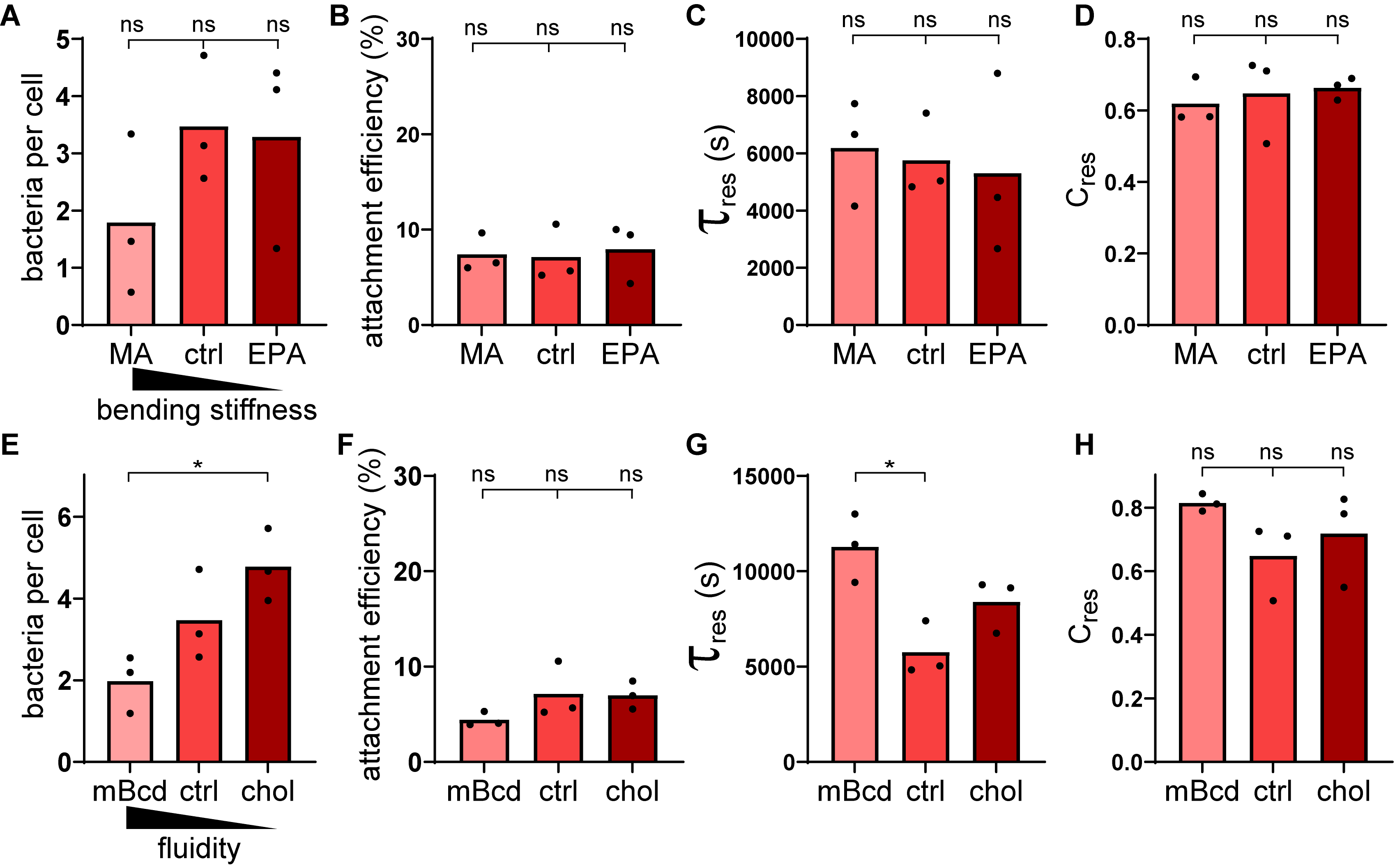

**Figure S6: Perturbations of mammalian membrane bending stiffness and fluidity have limited effect on bacterial attachment dynamics.**

(A-D) HeLa GFP were pre-cultured a saturated fatty acid, margaric acid (MA), or of a poly-unsaturated fatty acid, eicosapentaenoic acid (EPA), prior to flowing *E. coli* VHH. (A) Final *E. coli* VHH count per HeLa cell does not correlate with polyunsaturated fatty acid content. (B) Bacterial attachment probability does not correlate with polyunsaturated fatty acid content. (C,D) Characteristic residence time 𝜏_res_ and it pre-exponential factor C_res_ do not correlate with polyunsaturated fatty acid content. (E-J) HeLa GFP were cultured in cholesterol-depleting condition (methyl-β-cyclodextrin, mBcd), regular medium (ctrl), or cholesterol-enriching condition (chol) prior to flowing *E. coli* VHH. (E) Final *E. coli* VHH count per HeLa cell slightly increases with cholesterol content in flow. (F) Bacterial attachment probability does not correlate with cholesterol content. (G-H) Characteristic residence time 𝜏_res_ and its pre-exponential factor C_res_ do not correlate with cholesterol content. Statistical tests: one-way ANOVA followed by Tukey post hoc test (* P<0.05).

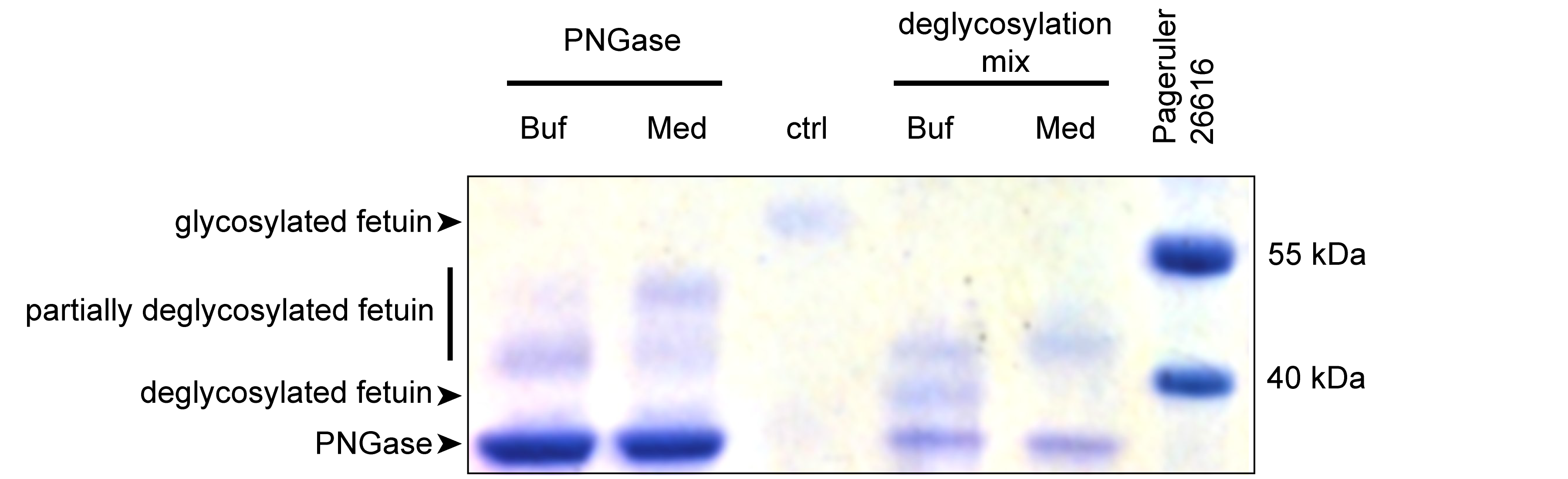

**Figure S7: Deglycosylation of a N- and O-glycosylated protein in GFP-displaying HeLa.**

SDS-PAGE gel showing 10 µg of fetuin incubated overnight at 37°C in 20uL with either only PNGase or the deglycosylation mix of enzymes in either the non-denaturing buffer provided by the supplier (Buf) or Fluorobrite medium (Med) supplemented with glutamax. The gel indicates partial deglycosylation of fetuin in mammalian cell culture conditions compared to untreated fetuin (ctrl) and compared to ideal buffer conditions.

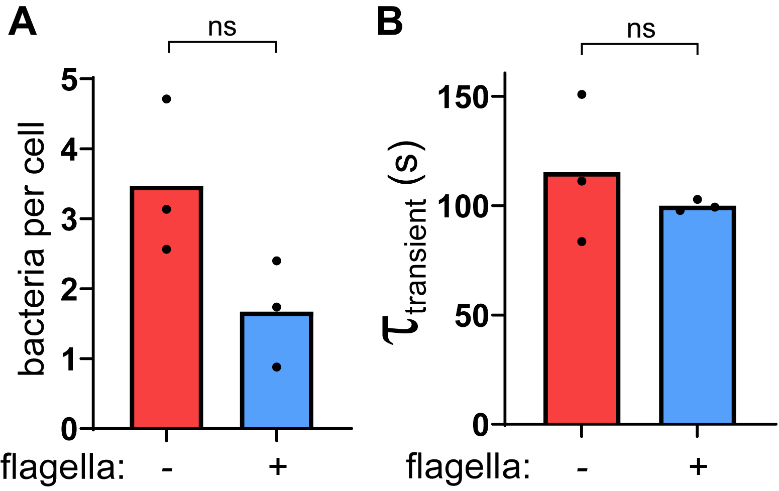

**Figure S8: Flagella does not affect overall attachment and unspecific interaction duration.**

(A) Final *E. coli* VHH count per HeLa cell is not significantly different in the presence or absence of flagella (“+” and “-”, respectively) in flow. (B) The average transient binding time is not affected by the presence of flagella. Statistical tests: two-tailed unpaired t-test.

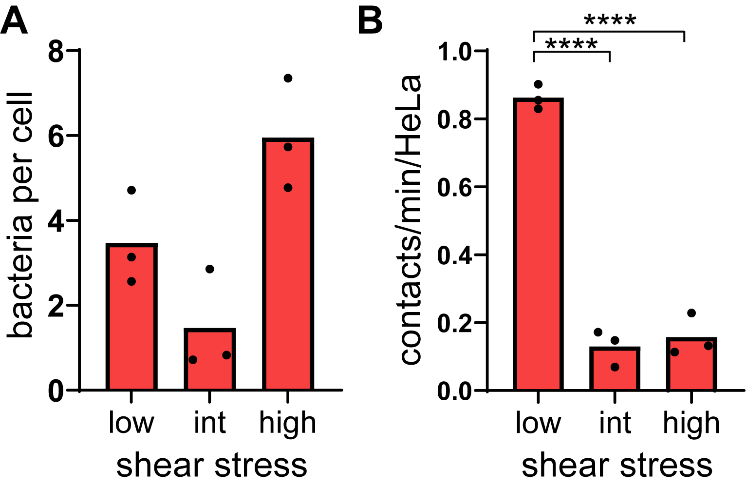

**Figure S9: The effects of flow on overall attachment and on contact frequency.**

(A) Final *E. coli* VHH count per HeLa cell is insensitive to flow. The data shows raw bacterial counts and is not normalized to the incoming flux of bacteria at different flow. (B) Strong flows decrease the contact frequency despite a higher number of bacteria crossing the channel. Statistical tests: one-way ANOVA and Tukey post hoc test (**** P<10^-4^).

**Supplementary Movies 1 (separate file): Live visualization of bacterial attachment to host cells.**

Microscopic visualizations of *E. coli* VHH adhesion to HeLa GFP. Maximum intensity projection of a 1-hour confocal microscopy time-lapse at 0.1 fps accelerated 100x. Overlay of bacteria (red) location of initial contact (circles) and their corresponding tracks (colored lines). Scale bar: 50 µm.

**Supplementary Movies 2 (separate file): High-speed visualization of bacterial attachment to host cells upon contact.**

Close-up on microscopic visualizations of *E. coli* VHH *E. coli* adhesion to HeLa GFP. Maximum intensity projection of 5-minutes confocal microscopy time-lapses at 1 fps accelerated 10x. Overlay of bacteria (red) location of initial contact (circles) and their corresponding tracks (colored lines).

**Supplementary Movie 3 (separate file): Attachment of *E. coli* VHH to GFP-coated coverslips.**

*E. coli* VHH in flow binding to the edge of a GFP-functionalized coverslip used as a substrate for a microfluidic channel. Maximum intensity projection of 5-minutes confocal microscopy time-lapse at 1 fps accelerated 10x. Scale bar: 50 µm.

**Supplementary Movie 4 (separate file): HeLa GFP cells actively pull bacteria towards their cell body.**

Maximum intensity projection of a confocal time-lapse experiment at 0.1 fps accelerated 100x of *E. coli* VHH (red), HeLa GFP (CD80) (green) and 3 mm/s flow. Scale bar: 10 µm.

**Supplementary Movie 5 (separate file): Bacteria sequester GFP upon attachment.**

Maximum intensity projection of 20-minutes epifluorescence microscopy time-lapse at 1 frame per minutes accelerated 60x. *E. coli* VHH were added on HeLa GFP under static conditions at an MOI of 200 for a couple of minutes and washed 3 times before imaging. Scale bar: 10 µm.

**Supplementary Movie 6 (separate file): Flagella promote unspecific transient binding.**

Maximum intensity projection of a confocal time-lapse experiment at 6 frames per minutes accelerated 100x of flagellated *E. coli* VHH (red), HeLa GFP (CD80) (green) and 3 mm/s flow. Scale bar: 10 µm.

**Supplementary methods**

**K_D(display)_ measurements**

To measure VHH affinity when displayed at the bacterium surface, serial dilution of GFP were performed while increasing the volume to avoid antigen depletion. For each concentration, 100 µL of bacteria induced with 250 ng/mL tetracycline overnight were washed with PBS and stained for two hours. For volumes below 50 mL, bacteria were pelleted and resuspended in 4% PFA in PBS. For volumes above 50 mL, bacteria were retrieved on 0.22 µm filters using 4% PFA in PBS. Bacteria were then imaged under a 1% agarose PBS pad. Wide field fluorescent pictures were taken at 100x and 1.5x lens magnification.

Using Fiji software, bacteria were detected using the mScarlet channel and the corresponding regions of interests were used to quantify mean GFP intensity for each bacterium. Prism software (Graphpad) was used to perform a non-linear fit of the mean GFP signal among bacteria on the field of views using the formula “One site – specific binding” Y = max * [GFP] / (K_D_ + [GFP]) and estimate K_Ddisplay_.

**HA tag staining**

Bacteria harboring a HA tag were washed with PBS and stained with anti-HA antibody conjugated with FITC (Abcam ab1208) at 10 µg/mL for 75 minutes in the dark, washed once with PBS and imaged under a 1% agarose PBS pad. Widefield fluorescent pictures were taken at 100x and 1.5x lens magnification.

**GFP uptake rate**

*E. coli* VHH were added under static conditions at a MOI of 200 for a couple of minutes and washed 3 times before widefield epifluorescence imaging. HeLa GFP captured 11 bacteria. Image segmentation performed using the red channel (*E.coli*) to quantify the local total GFP signal around bacteria over time.

**Supplementary table 1: Plasmid cloning strategy**

| Name | Description, cloning | Sources |
| --- | --- | --- |
| pDSG323 | Empty display control, Kan^R^  Tet-inducible truncated intimin construct | Glass et al ^1^, Addgene 115594 |
| pDSG339 | VHH anti GFP display. Kan^R^ , K_D_ = 0.59 nM  Tet-inducible VHH display based on truncated intimin | Glass et al^1^ |
| PeGFP_GPI | pEGFP-N1 - preproinsulinSP eGFP linker DAF GPI  eGFP display for mammalian cells , anchoring motif from CD55, preproinsulin secretion peptide | Ricci et al ^2^  Generous gift from Prof. Van der Goot, EPFL |
| pXP145 | Constituve eGFP (N105Y, E125V,Y146F) display based on C-terminal CD80 anchor, Amp^R^, Neo^R^.  Digestion and ligation of the following:  Backbone: pCDNA3* XbaI HindIII  Insert: pENTR_SignalPeptide-GFP(N105Y/E124V/Y145F-superfastFolding** mutation)_mCD80TransMembrane XbaI HindIII | *Invitrogen  **Generous gift form Prof. Joerg Huelsken, EPFL  This study |
| pXP226 | eGFP in pET28a for recombinant expression  Gibson assembly with the following PCR products:  pET28a* oXP546 oXP547  pUCBB** oXP333 oXP334 | *EMD Biosciences  **Vick et al ^3^,  Addgene 32548  This study |
| pXP327 | Dox-inducible eGFP (N105Y, E125V,Y146F) display based on CD80 anchor on lentivector, Amp^R^, Neo^R^  Digestion and ligation of the following:  Backbone: pCW57-RFP-P2A-MCS* EcoRI BamHI  Insert: pXP145 first amplified with oXP799 oXP800 then EcoRI BamHI | *Barger et al^4^, Addgene 78933  This study |
| pXP340 | Dox-inducible eGFP (N105Y, E185V,Y206F) display based on CD80 anchor on lentivector, Amp^R^, Neo^R^  Digestion and ligation of the following:  Backbone: pRRLSIN.cPPT.GFP.WPRE* SacII NotI  Insert: pXP327 SacII NotI | *Trono lab, EPFL, unpublished  Addgene 12252  This study |
| pXP383 | VHH anti GFP display. Kan^R^ , K_D_ = 0.59 nM  Tet-inducible VHH display based on truncated intimin + HA tag.  Gibson assembly with the following PCR products:  pDSG339 oXP912 oXP926  pDSG339 oXP528 oXP913 | This study |
| pXP384 | Low affinity VHH anti GFP display. Kan^R^ , K_D_ = 16 nM  Tet-inducible VHH display based on truncated intimin + HA tag.  Gibson assembly with the following PCR products:  pDSG339* oXP914 oXP915  LaG02* oXP916 oXP917 | *Fridy et al ^5^  This study |
| pXP388 | High affinity VHH anti GFP display. Kan^R^ , K_D_ = 20 pM  Tet-inducible VHH display based on truncated intimin + HA tag.  Gibson assembly with the following PCR products:  pDSG339* oXP914 oXP915  LaG94-10* oXP924 oXP925 | *Fridy et al ^5^  This study |
| pZA002 | pGRG36 j23119_mScarlet  Constitutive synthetic promoter driving the expression of mScarlet in Tn7 vector.  Digestion and ligation of the following:  Backbone: pGRG36* PacI XhoI  Insert: synthetized j23119**_mScarlet PacI XhoI | *McKenzie et al ^6^  Addgene 16666  **parts.igem.org/  Part:BBa_J23119  This study |
| SpyTag003-mKate2 | Expresses SpyTag003-mKate2 (a far-red fluorescent protein) in bacterial cytoplasm | Keeble et al, ^7^  Addgene 133452 |

**Supplementary table 2: Primer table**

| oXP333 | ATGGTGAGCAAGGGCGAG |
| --- | --- |
| oXP334 | TCACTTGTACAGCTCGTCC |
| oXP528 | ttaccaatgcttaatcagtgagg |
| oXP546 | ATGGACGAGCTGTACAAGTGAGATCCGGCTGCT |
| oXP547 | CTCGCCCTTGCTCACCATGCTGCTGTGATGATGATG |
| oXP799 | agcgaattcgccaccatggactcc |
| oXP800 | cccggatccctaaaggaagacggtctgttc |
| oXP851 | atcaggcaatttggcgttgccgtcagtctcagttaatcaggttacaacgagtgtaggctggagctgcttc |
| oXP852 | agaagcgtagccgtaatcggattattcgcgagccatcgactcattcagatggtccatatgaatatcctccttagttcc |
| oXP912 | TACCCGTATGATGTTCCCGACTATGCCatggctcaggtgcagctg |
| oXP913 | GGCATAGTCGGGAACATCATACGGGTAtctagtCGCACCATCAAAAAATATAAC |
| oXP914 | TAATAAtactagtagcggccgc |
| oXP915 | GGCATAGTCGGGAACATCATAC |
| oXP916 | GTTCCCGACTATGCCATGGCCCAAGTTCAGCT |
| oXP917 | cgctactagtaTTATTATACAGTAACCTGTGTTCCCTG |
| oXP924 | GTTCCCGACTATGCCATGGCTCAAGTCCAGCTTG |
| oXP925 | gctactagtaTTATTAACTGACGGTCACCTGC |
| oXP926 | gagtcaggcaactatggatgaac |
